# Argonaute-mediated system for supersensitive and multiplexed detection of rare mutations

**DOI:** 10.1101/803841

**Authors:** Qian Liu, Xiang Guo, Guanhua Xun, Zhonglei Li, Yuesheng Chong, Litao Yang, Hongxia Wang, Fengchun Zhang, Shukun Luo, Zixin Deng, Kai Li, Yan Feng

## Abstract

The ability to detect rare mutations has revolutionized diagnosis and monitoring of tumors, but is limited by the shortage of sensitive, cost-effective and high coverage methods for identification of extremely low abundant mutations. Here, we establish a single-tube multiplex PCR system by employing thermophilic Argonaute-derived DNA-guided nuclease for highly efficient rare mutation detection, referred to as A-Star (Argonaute-directed specific target enrichment and detection), that combines the selective cleavage of the wild type DNA in the DNA denaturation step and the followed amplification of mutant DNA during PCR. A-Star enables easy detection and quantitation of rare mutations originally as low as 0.01% in allele frequency with a ⩾ 5500-fold efficiency. We also demonstrate the feasibility of A-Star for detecting oncogenic mutations in complex biological systems such as solid tumors tissues and blood samples. Remarkably, A-Star could achieve the detection of multiple oncogenic genes simultaneously through a simple single-tube reaction. Taken together, our work illustrates a supersensitive and rapid nucleic acid detection system, thereby extending the utility for both research and therapeutic applications.

---

The detection of rare polymorphic alleles is becoming increasingly relevant for the early diagnosis and monitoring of a variety of tumors^1,2^. The clinical detection methods for rare mutations, including single nucleotide variations (SNVs) and insertion/deletion (indel) mutations, that are currently available involve either sequencing or PCR^3, 4^. Compared with authentication by sequencing DNA fragments, allele-specific diagnostic PCR is not only simpler and timesaving but also practical and effective^5-7^. Therefore, the detection of extremely rare variant alleles within a complex mixture of DNA molecules has attracted increasing attention aimed at solving the technical problems associated with the strict requirements for both a precise single-nucleotide resolution and simple multiplex detection, especially in the detection of circulating tumor DNA in the circulated tumor DNA of patients^8-10^.

Recently, a variety of rare mutation detection methods based on endonucleases with sequence-specific catalytic capabilities^11,12^, particularly the enzymes of the CRISPR/Cas systems, have been developed^13-15^. These strategies are primarily based on Cas9 cleavage activity^16,17^ or the collateral cleavage activity of Cas12 or Cas13a effector^18,19^. Once combined with the pre-enrichment of recombinase polymerase amplification (RPA), 5% SNV allele can be achieved in the Cas12/13-enzymes^20-22^. However, the major limitations of these approaches are different cutting preferences needed for the vicinity of target sequence cleavage and the designed reporter system^20-24^, which restricts the utility of these methods for the detection of a wide spectrum of target genes, also poses a great challenge in the multiplexed detection that needed tedious screening for orthogonal CRISPR enzymes in targeting multiple genomic loci simultaneously^24^. Additionally, the system including the synthesized guide RNA, the target RNA converted from the pre-amplified ctDNA by T7 transcription, and Cas enzymes reaction causes the quite high-cost and technical inconvenience.

One potential means of circumventing these limitations is to take advantage of the unique properties of thermophilic Argonaute (Ago) proteins, which do not require a prerecognition motif and instead specifically cleave target nucleic acids via the base-pairing with the small guide nucleic acids^25,26^, such as reported from *Pyrococcus furiosus, Thermus thermophiles, Methanocaldococcus jannaschii* and *Methanocaldococcus fervens, etc*^27-30^. The revealed capacity of Agos from hyperthermophile *Pyrococcus furiosus* (*Pf*Ago), which had been biochemistry characterized and designed as a programmable DNA-guided artificial restriction enzyme^27,31^, caught our great attention with functioning precise cleavage of the target DNA (tDNA) between positions 10 and 11 of the pairing guide DNA (gDNA) at temperatures as high as 95 °C. We assume that the powerful catalytic ability of *Pf*Ago for the single strand DNA (ssDNA) may allow it to cleave the unwinding double strands DNA (dsDNA) during the first denaturing step of PCR. A deep understanding and diligent exploration of *Pf*Ago, through how gDNA-directed Ago discriminates the wild-type (WT) DNA and rare variant allele precisely, how to combine *Pf*Ago into a PCR reaction to enrich rare mutations efficiently, and whether *Pf*Ago can deal with multiplex detection rapidly, etc., may facilitate the development of new protocols that enable the rare mutation detection with great precision and efficiency.

Here, we report the establishment of A-Star (<u>A</u>go-directed <u>S</u>pecific <u>Tar</u>get enrichment and detection), a simple but supersensitive single-tube PCR system that takes advantage of *Pf*Ago to specifically cleave WT sequences in the denaturing step of PCR, leading to progressive and rapid enrichment of rare variant-containing alleles during the PCR process. As a proof of concept, the compatibility of *Pf*Ago with PCR is evaluated by introducing the well-designed gDNA directly in a single tube, which demonstrate the highly efficiency by orders of magnitude compared with a divided system between the WT cleavage and variant alleles amplification. The mocked cell-free DNAs (cfDNAs) and clinical human samples are tested for target oncogene enrichment. Furthermore, a multiplexed variant alleles system for one-pot reliable detection with single-digit copies is demonstrated by multi-gDNA directed *Pf*Ago system in the orthogonal manner. Our investigation indicates that the A-Star method we have developed provides a novel, supersensitive, rapid and inexpensive way in rare variant alleles detection, which exhibits great potential from applications in basic research to potential new diagnostic tools and clinical utility.

## Results

### Design of A-Star platform

We reasoned that *Pf*Ago can be employed in variant alleles enrichment and detection by coupling PCR analysis in a single-tube. A flow chart of the A-Star is shown in Fig. 1. During the DNA denaturation step at 94 °C, the designed and optimized gDNAs extend the precise complementation of WT ssDNA but not variant alleles, consequently to direct the *Pf*Ago-specific cleavage of unwinding dsDNA and enabling the elimination of a large fraction of the WT background. The subsequent annealing and strand-extension steps initiated by routine PCR primers and DNA polymerase favor the amplification of the spared variant alleles template rather than the cleaved WT fragments. The catalytic elimination of WT accompanied by the exponential amplification of the variant alleles in each cycle of PCR led to specific enrichment of the mutant target in an efficient and simple performance. Coupling the A-Star approach with TaqMan real-time quantitative PCR (TaqMan PCR) or Sanger sequencing allows the detection of the enriched rare variant alleles (full details are presented in the Methods).

**Fig. 1.**
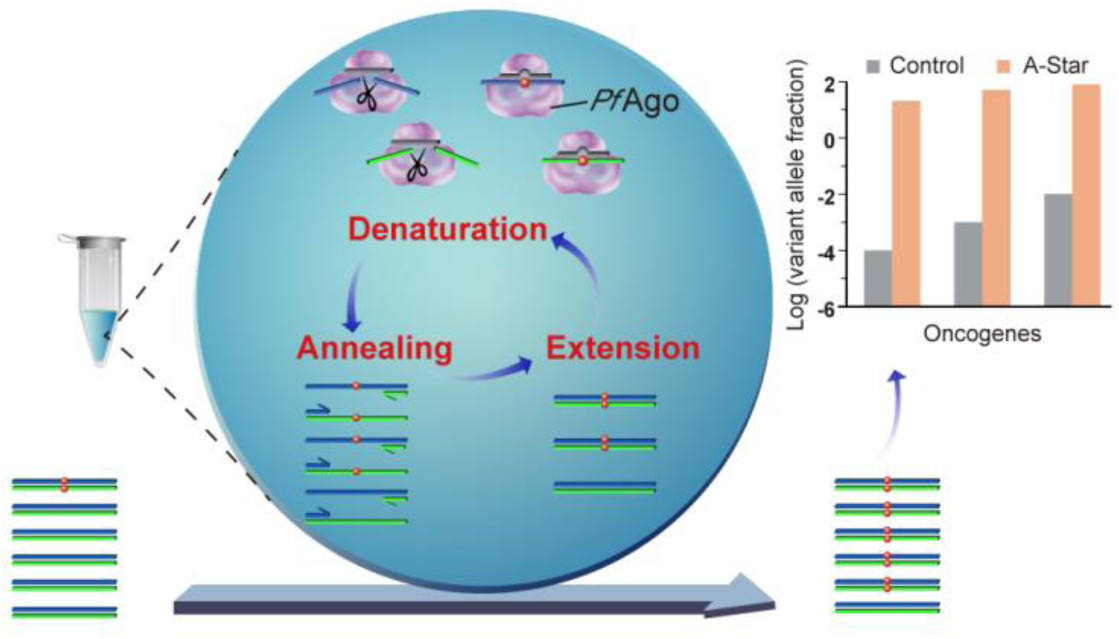
Schematic representation of the A-Star approach for rare mutation enrichment. Black lines represent gDNA, whereas blue and green lines represent forward or reverse strand of target dsDNA; Red balls indicate the mutated nucleotide of SNV target; the lines with arrows represent the primers.

We first tested the feasibility of thermophilic Ago used for PCR by examining the temperature dependence of *Pf*Ago cleavage activity against ssDNA targets. When the *Pf*Ago-gDNA complex was incubated with a synthesized 60 nt *KRAS* ssDNA fragment for 15 minutes, which is longer than the total cumulative time of the DNA denaturing step in PCR, we observed high cleavage activities at temperatures ranging from 80 to 97 °C but not below 70 °C (Supplementary Fig. 1), indicating that *Pf*Ago catalysis is not only compatible with the DNA denaturing step of PCR at 94 °C but also favorable for continuous WT cleavage during the PCR thermal cycling. To avoid nonspecific extension initiated by the gDNA in subsequent PCR amplification for both the WT and rare variant alleles, we modified each gDNA by adding an extra 3’-phosphate and verified its effectiveness by pre-PCR (Supplementary Fig. 2). Taken together, the results indicated that *Pf*Ago cleavage and PCR amplification are compatible in a single reaction used with a 16 nt gDNA in which both the 5’- and 3’-termini are modified by phosphate groups.

### gDNA design enables precise discrimination between the WT and SNV

To address the precise cleavage of the WT alleles, the crucial concern in A-Star involves the design and selection of the gDNAs for the precise discrimination of tDNAs at single nucleotide resolution. We optimized the gDNA using the synthesized *KRAS* ssDNA fragment of the WT or G12D allele (a well-studied oncogene, c.35G>A) as a representative cleavage target. Considering the potential effect of mismatch position and nucleotide type on DNA cleavage efficiency^28^, we applied a systematic design for selecting the best gDNA hit for the highly specific cleavage of the WT sequence. When 16 nt gDNA sequences perfectly matching the WT sequence were incubated with *Pf*Ago and *KRAS* ssDNA at 94 °C for 15 min, both WT *KRAS* and mutant *KRAS* containing the SNV encoding a G12D substitution were cleaved indifferently (Supplementary Fig. 3). This result suggests that the single-nucleotide difference in the gDNA for the WT and SNV alleles is not sufficient for discriminating DNA cleavage.

To improve the discrimination between the WT and SNV tDNA sequences, we introduced an additional mismatched nucleotide of the gDNA at positions around its SNV pairing site and then evaluated the effect on cleavage efficiency. When a mismatch was introduced at position −4, −1, or +1 of the gDNA, corresponding to gDNA gM7, gM10 and gM11, respectively, *Pf*Ago preferred to cleave the WT *KRAS* ssDNA, resulting in the lowest cleavage efficiency of SNV allele (Fig. 2a and Supplementary Fig. 4). The same phenomenon was observed on the WT and E545K mutation of *PIK3CA* (data not show). Due to the importance of the flanking positions of the cutting site of the gDNA, we focused on double, continuous mismatches at positions −1 and +1 of the SNV site for further gDNA optimization. We also observed that the mismatched nucleotide identity of the gDNA affects the discrimination performance of *Pf*Ago, but with a high dependency on the target sequence, not the specific nucleotides (G/C or A/T) (Supplementary Fig. 5a-d and Supplementary Fig. 6a-d). Collectively, these results indicated that introducing mismatch on gDNA at corresponding cutting sites to constitute continuous double mismatches with SNV sequence can direct the preferential cleavage of WT sequences, thereby allowing SNV selection. Considering the complexity of the gene sequences, we generally expand the gDNA candidate spectrum for the best hit from two gDNA groups with mismatched nucleotides at positions (−1, 0) and (0, +1) for the SNV allele in the following work to discriminate WT and rare SNV alleles.

**Fig. 2.**
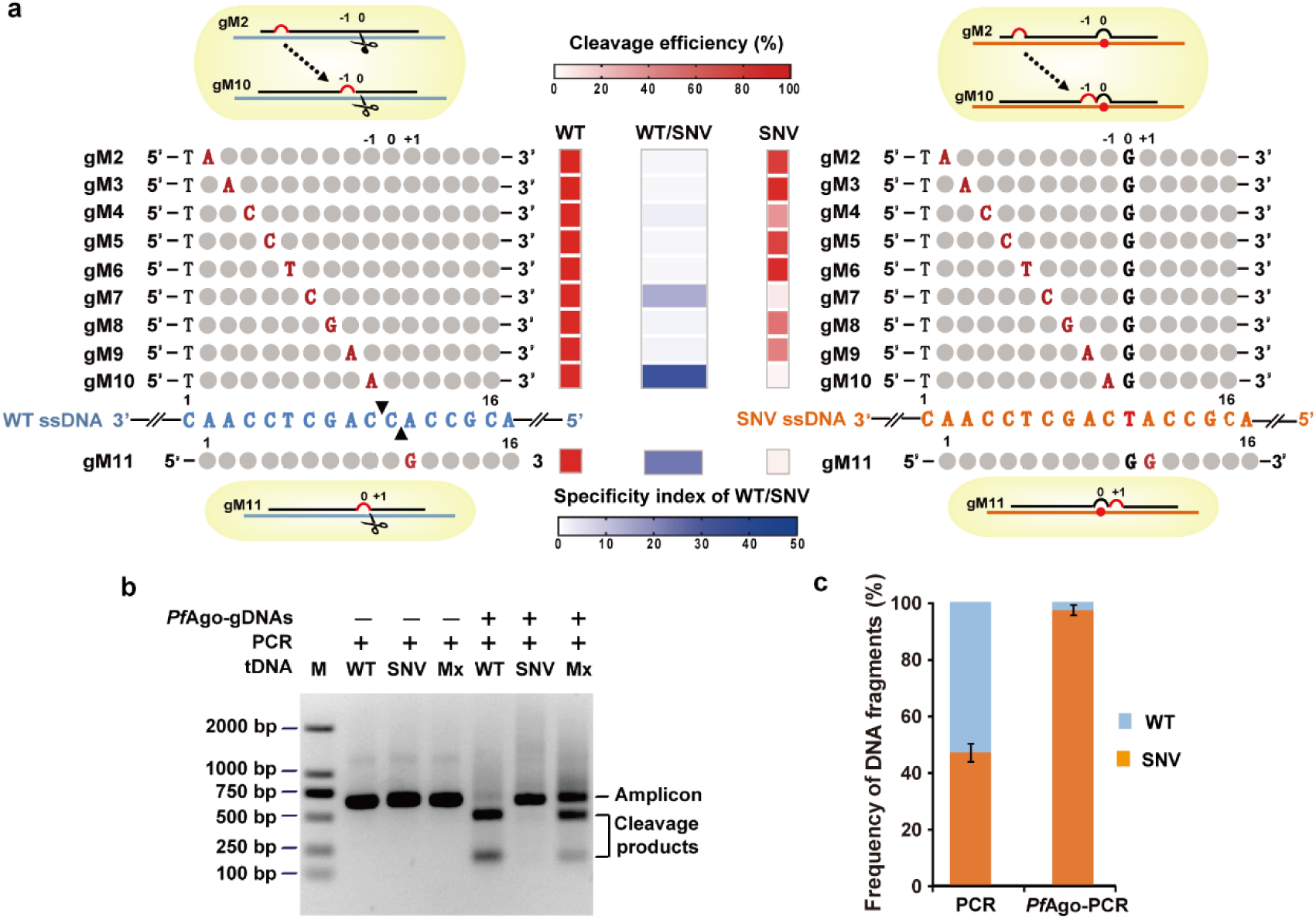
Discriminated DNA cleavage by designed gDNAs and its compatibility with PCR in a single-tube reaction for SNV enrichment. **a** Schematic representation of designed gDNAs for *KRAS* G12D ssDNA cleavage. For the designed gDNAs, the introduced mismatched nucleotides are highlighted in red, and the nucleotides corresponding to the SNV site is highlighted in black and labeled as “0” site. The black triangles on the WT ssDNA represent the specific cleavage sites. The cleavage efficiency was quantified by measuring the electropherogram of the cleavage products with the ssDNA target of *KRAS* G12D WT or SNV, respectively, as shown in the red heatmap. The specificity index of the WT/SNV is calculated as the ratio of the cleavage efficiency of WT to that of SNV, as shown in the blue heatmap. **b** Gel electrophoresis analysis for *Pf*Ago-PCR coupled reaction to cleave WT sequences and enrich SNVs. Mx: mixture of WT and SNV target at equal mole. **c** TaqMan PCR evaluation of the enrichment results tested with a mixture of WT and SNV target at equal mole by *Pf*Ago-PCR coupled reaction. Error bars represent the mean ± s.d., n = 3.

After optimizing the gDNA candidates for each single strand of the WT sequence, we further determined the recombination effect of the paired gDNAs on the discriminating cleavage of a dsDNA target to completely eliminate the WT in the PCR system. When we used a pair of best hit gDNAs, the SNV variants of *KRAS* G12D and *PIK3CA* E545K remained uncleaved, whereas their corresponding WT counterparts were almost completely cleaved by incubation with *Pf*Ago at 94 °C for 15 min (Supplementary Fig. 5e-f and Supplementary Fig. 6e-f). When a pair of semi-best hit gDNAs in which one best gDNA was mixed with a poorer hit gDNA was used, we did not observe clear discriminating cleavage of the dsDNA target. This result verified that one pair of the best hit gDNAs could direct *Pf*Ago to carry out discriminating cleavage of each individual strand of the unwinding dsDNA. Additionally, considering that the indel mutation is quite popular in oncogenes, we asked whether a pair of gDNAs matched to the WT sequence could discriminate between the WT and mutant dsDNA target. Taking *EGFR* and its 15-nt deletion mutant as an experimental case, it was confirmed that *Pf*Ago can efficiently cleave WT dsDNA with a minimal effect on the mutant allele at 95 °C (Supplementary Fig. 7). Taken together, these results demonstrated that *Pf*Ago is capable of effectively discriminating mutated dsDNA from WT alleles when directed by a pair of gDNAs that each contains two consecutive nucleotides mismatched with the SNV templates at position +1 or −1.

### A-Star specifically enriches rare SNV in a single-tube

We next asked whether gDNA-directed *Pf*Ago targeting could be combined with a routine PCR amplification program in a single reaction tube. We tested the effect of *Pf*Ago on the efficient cleavage of tDNA by using the *KRAS* G12D WT and SNV alleles, respectively, in the presence of a pair of best hits gDNAs. PCR was performed with amplification cycles of 94 °C for 30 s, 55 °C for 30 s, and 72 °C for 20 s after pre-incubation, as suggested by the commercial protocol. Gel electrophoresis provides direct proof of discriminating cleavage since the products from WT are almost completely digested into two small fragments, whereas SNV alleles are enriched as single, full-length amplicon (Fig. 2b). When a mixture of WT and SNV alleles at a ratio of 1:1 was used as a template, three bands appeared including the full-length amplicon and two small fragments. We further evaluated SNV allele enrichment quantitatively by TaqMan PCR. The results showed that more than 90% of the amplicons in the mixture was analyzed as the SNV alleles in the presence of *Pf*Ago-gDNAs, and approximately 50% of the amplicons were analyzed as the SNV alleles by routine PCR treatment (Fig. 2c). Sanger sequencing also provided support for the enrichment of SNV alleles, since the guanidine nucleotide peak from WT became a major peak, differing from the results for the PCR-treated sample showing two distinct peaks at equivalent level (Supplementary Fig. 8). Collectively, the results indicated that the PCR approach combined with the use of gDNAs-directed *Pf*Ago could led to SNV enrichment by eliminating WT alleles in a single tube.

Seeking to increase the detection sensitivity for rare SNV alleles using a 1% variant allele fraction (VAF) of *KRAS* G12D, we systematically titrated the *Pf*Ago amount, the molar ratio between *Pf*Ago and gDNA, and the PCR thermal cycling conditions. We found that the *Pf*Ago concentration greatly affected SNV enrichment (Supplementary Fig. 9). As the *Pf*Ago concentration was increased to 30 nM with a ratio of 3:1 between *Pf*Ago and tDNA, the majority of WT alleles could be removed, leading to efficient rare SNV enrichment; when the concentration was over 30 nM and the molar ratio reached 4:1 or 5:1, the SNV enrichment efficiency decreased, which might have been caused by the strong binding ability of *Pf*Ago toward unspecific DNA and thus reduced the efficiency of tDNA concentration for catalytic cleavage. We also found that an increased ratio of 20:1 between gDNA and *Pf*Ago could enhance SNV enrichment (Supplementary Fig. 10), implying that gDNA with a high concentration could competitively bind *Pf*Ago and release the bound tDNA for further specific cleavage. Moreover, during the PCR procedure, 25∼30 thermal cycles could efficiently enrich the rare SNV alleles (Supplementary Fig. 11). Taken together, the results showed that the optimal conditions were 30 nM *Pf*Ago with a *Pf*Ago:gDNA ratio of 1:20, with a 25-cycle PCR program.

### A-Star performs a super enrichment ability for rare SNV

We further evaluated the limit of detection (LOD) of A-Star according to the differential VAFs of *KRAS* G12D at 0.01%, 0.1%, and 1% levels. A-Star could enrich SNV alleles efficiently, ranging from 55 to 75% as detected by TaqMan PCR (Fig. 3a), although the initial SNV VAFs varied by several orders of magnitude. The results showed that the samples at VAF of 0.1% and 1% were enriched to over 60%, responding to 600 and 75 fold increase, respectively. Remarkably, even for a sample with a VAF as low as 0.01%, the SNV products could be enriched for over 5500-fold (Fig. 3b) and were readily detected by Sanger sequencing (Supplementary Fig. 12). Similar results were obtained at a VAF of 0.01% for samples containing a *PIK3CA* SNV or an *EGFR* indel mutation (Supplementary Fig. 13). To prove the advantage of A-Star over the uncoupled *Pf*Ago PCR reaction for the efficient enrichment of rare variant alleles, we compared A-Star with uncoupled *Pf*Ago PCR of 0.1% and 1% K*RAS* G12D, in which the latter reaction was set up as a two-step reaction, firstly, 15 min incubation with the *Pf*Ago-gDNA complex at 94 °C for WT cleavage and then 25 cycles PCR for the purified DNA products without the *Pf*Ago disturbance. We observed a marked difference in the enrichment efficiency of rare mutations between our single-tube *Pf*Ago-PCR and the uncoupled process. The A-Star technique increased the percentage of *KRAS* G12D SNV sequences in the final PCR products from 1% to 75%, which is more efficient than the uncoupled strategy which just increases to 6% (Supporting Information, Figure S14 and S15). It demonstrates that the simultaneous gDNA-directed *Pf*Ago cleavage of WT during PCR cycles in a single tube could enhance SNV allele enrichment significantly compared to uncoupled enzyme catalytic cleavage and PCR. In addition, we evaluated the mean cycle threshold (Ct) values of the samples with different VAFs and found an obvious linear relationship (Fig. 3c), indicating a high potential for quantitative analysis of the rare mutations on oncogenes in the early stage diagnosis or monitoring of oncogenes.

**Fig. 3.**
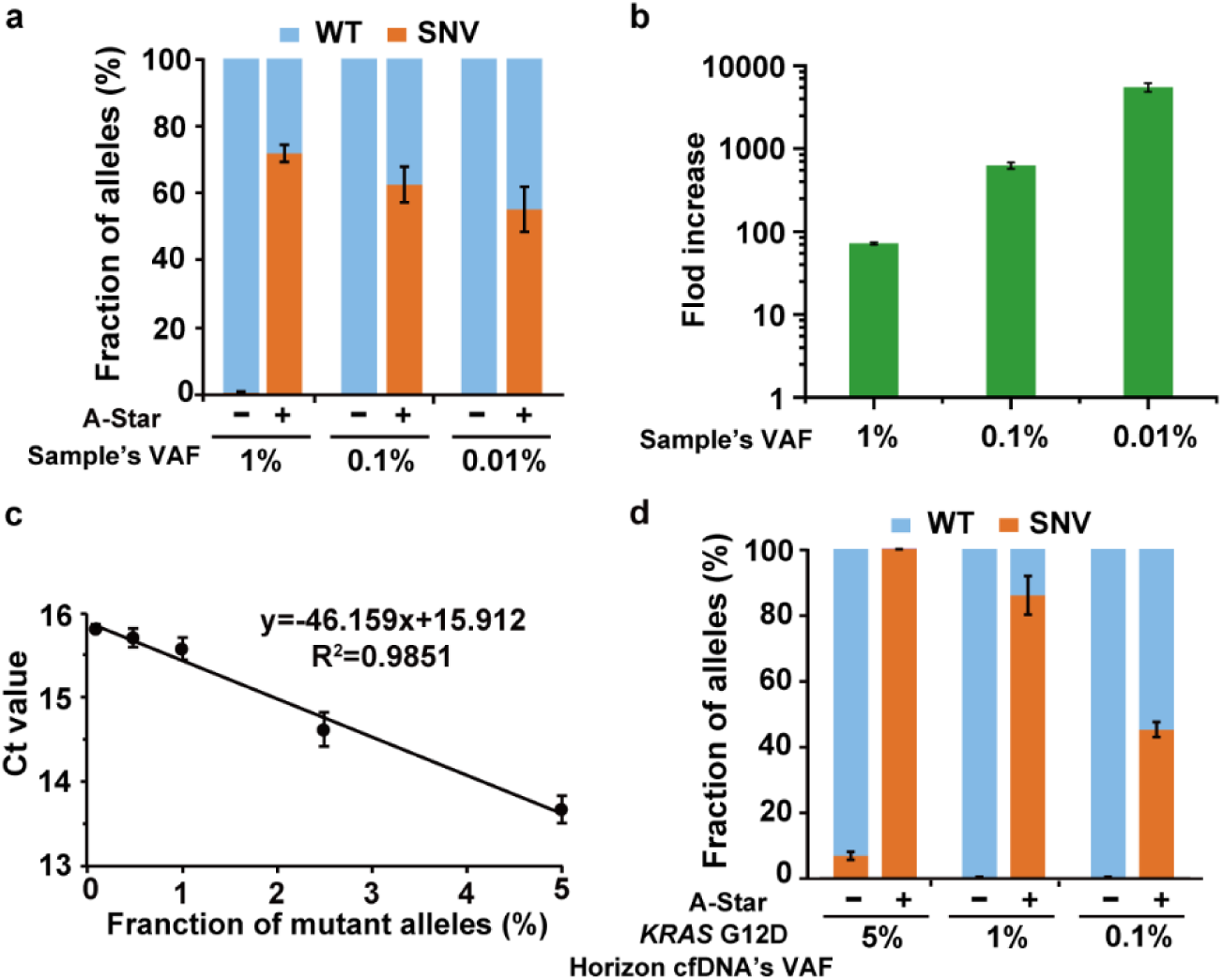
Detection sensitivity of A-Star for rare SNV enrichment. **a** Evaluation of enrichment efficiency of A-Star with *KRAS* G12D samples of varying VAFs by TaqMan PCR. **b** The values of fold increase were calculated from the data of (a). **c** The linear correlation of the threshold cycle (Ct) value with the VAFs tested by *KRAS* G12D samples and quantified by TaqMan PCR assays. **d** Evaluation of the A-Star enrichment results for *KRAS* G12D Horizon cfDNA standard samples of 0.1%, 1%, and 5% VAFs. Error bars represent the mean ± s.d., n = 3.

To explore the clinical utility of A-Star, we tested the LOD of A-Star using a set of commercial cfDNA standards containing genomic DNAs with three available VAFs of 0.1%, 1%, and 5% for *KRAS* G12D, *PIK3CA* E545K and the *EGFR* indel. Since the low amount of cfDNA in mock samples was not directly detectable using the A-Star method (data not shown here), we added 30 cycles of pre-PCR amplification to increase the amount of cfDNA for the WT and variant alleles. The inclusion of this step enabled the enrichment and detection of SNVs from samples containing only 33 ng of cfDNA with a VAF of as low as 0.1% for the variants (Fig. 3d and Supplementary Fig.16), that equals to single-digit copies of rare mutation. The sensitivity and specificity of A-Star on the mock cfDNA samples emphasize its potential for analyzing samples with high-background genomic DNA.

### A-Star can be used for clinic analysis of tissue and blood samples

We also tested A-Star on DNA extracted from tissue and blood samples from patients with different cancer types who carried the *KRAS* G12D SNV identified by deep sequencing (Fig. 4a). After A-Star processing, the SNV was successfully detected by TaqMan PCR and Sanger sequencing (Fig. 4b and Supplementary Fig. 17). The VAFs of tissue samples processed by A-Star directly increased up to 60-90% from the original VAFs of less than 20%. In particular, the LUAD sample with an extremely low VAF (below 1%) was enriched by more than 50%, representing over a 60-fold improvement. For blood samples, we performed a pre-amplification step to increase the initial template amount and then the A-Star procedure. The VAFs increased up to 40-85% from the original VAFs of 2-20%. The above results prove that A-Star offers impressive specificity and sensitivity for rare SNVs detection in clinical application.

**Fig. 4.**
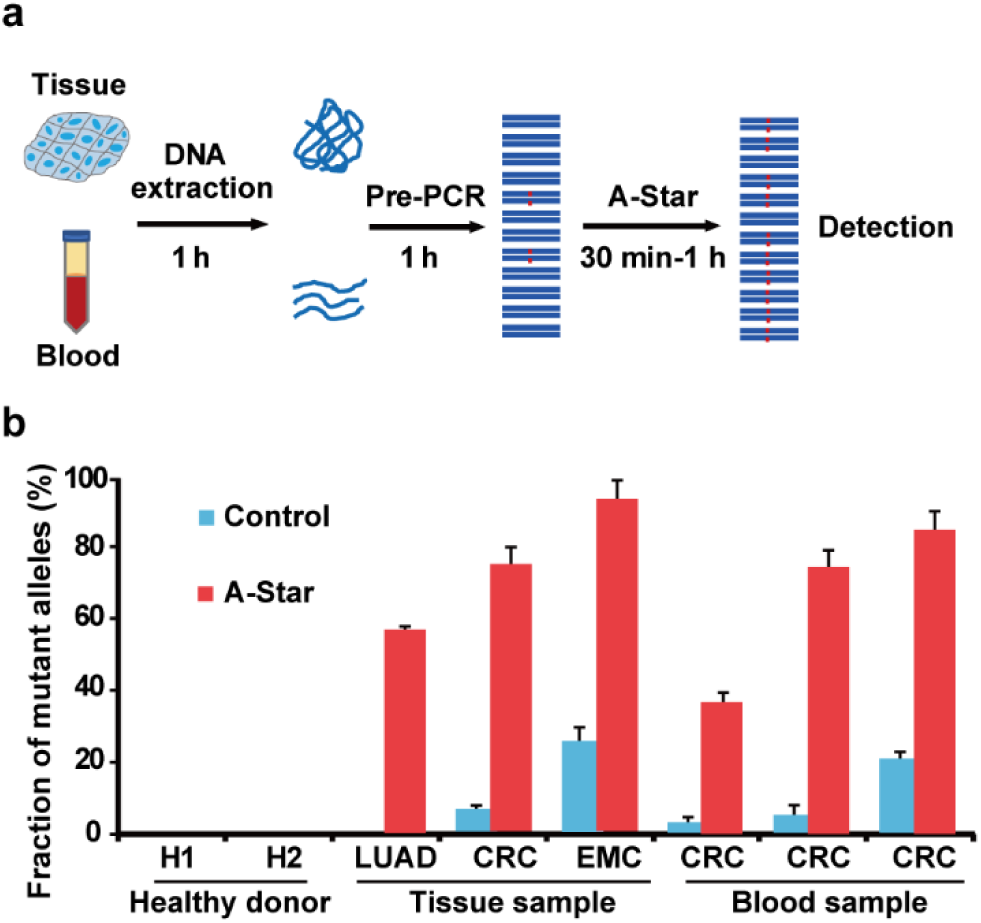
Analysis of A-Star in human clinic samples. **a** Flow chart of the A-Star procedure for the identification of cancer-associated SNVs in human samples. **b** Evaluation of A-Star by TaqMan PCR tested with clinical samples from diverse cancer types, as exemplified by *KRAS* G12D-containing samples from patients with lung adenocarcinoma (LUAD), colorectal cancer (CRC), or endometrial cancer (EMC). The control reactions did not contain gDNA pairs. Error bars represent the mean ± s.d., n = 3.

### Multiplexed detection of oncogenes

Given that Ago’s precise recognition capacity is mediated by gDNAs, we envisioned that A-Star in combination with properly multiple designed gDNAs might be suitable for the multiplex mutation detection (Fig. 5a). First, we tested the duplex enrichment efficiency by a different combination of variant allele targets among *KRAS* G12D and *PIK3CA* E545K and a deletion mutation for *EGFR* with a 1% VAF. The samples were mixed with all the required primers, designed gDNAs (Supplementary Fig. 18), and *Pf*Ago in one tube. The degree of enrichment of all targets analyzed by subsequent TaqMan PCR analysis was of a similar magnitude to the values detected for nonmultiplex A-Star assays (Fig. 5b and Supplementary Fig.19). Under triplex enrichment in one tube, all three targets were more than 40-fold enriched, implying orthogonal activity of *Pf*Ago when supplied with multiple gDNA pairs.

**Fig. 5.**
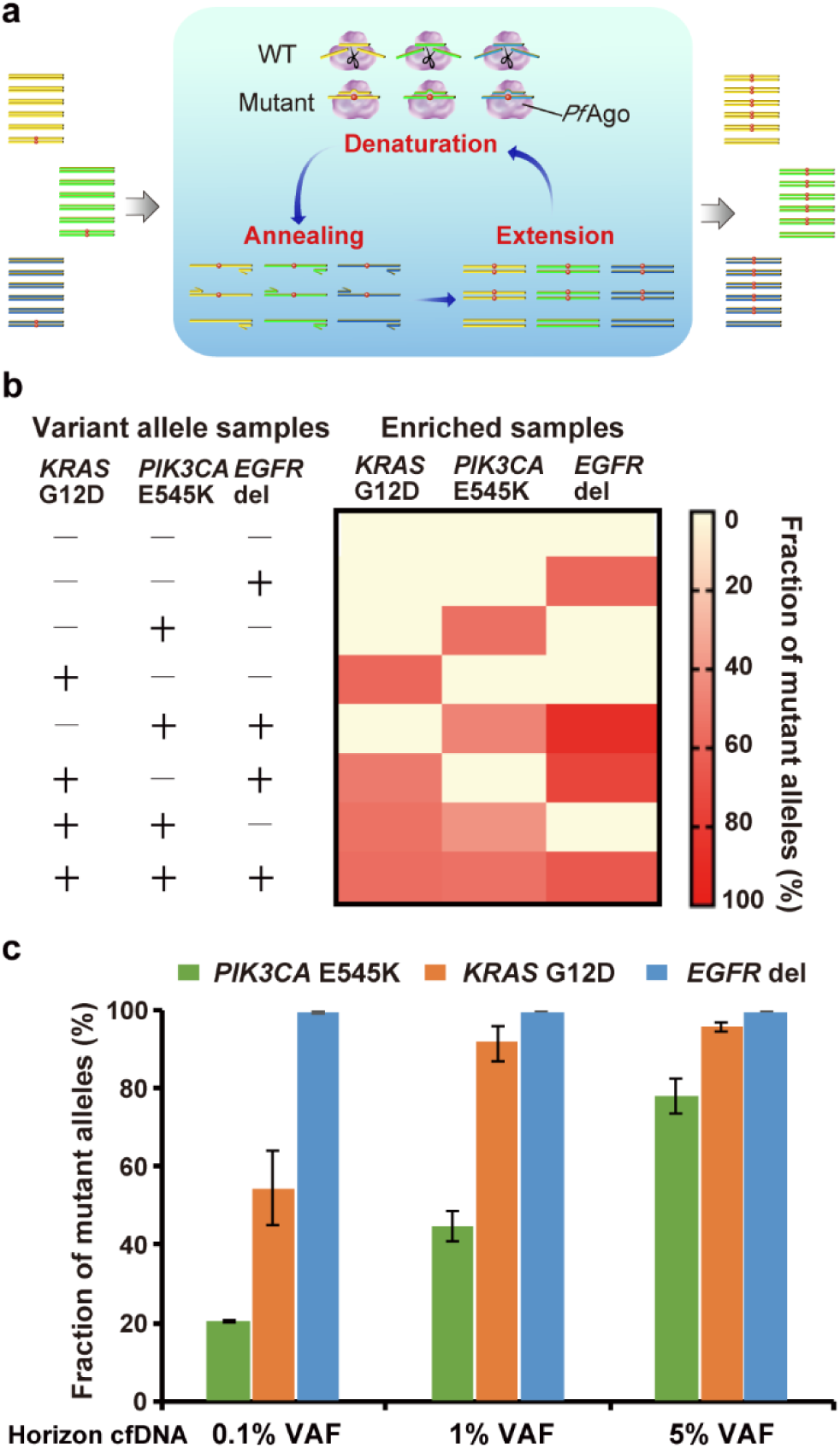
Multiplex detection of A-Star. **a** Schematic representation of multiplex mutation detection in a single-tube reaction containing multiple pairs of primers and gDNAs. **b** Triplex detection of A-Star by 1% VAF samples contacting *KRAS* G12D, *PIK3CA* E545K, and *EGFR* del target. **c** A-Star triplex enrichment of *KRAS* G12D, *PIK3CA* E545K, and *EGFR* del using standard cfDNA samples with different VAFs of 0.1%, 1%, and 5%. The enrichment output was detected using TaqMan PCR. Error bars represent the mean ± s.d., n = 3 for each variant.

To mimic clinical samples with a complicated DNA background, we also tested triplex A-Star on mocked cfDNA standards with three available VAFs (0.1%, 1%, and 5%) following a similar multiplex procedure as above but with an initial pre-PCR amplification step. The quantitative analysis revealed 50-90% of variant alleles for each of *KRAS* and *PIK3CA*, with a range of originally 0.1% to 5% SNV alleles, except for 0.1% PIK3CA (Fig. 5c and Supplementary Fig. 20). For 0.1% *PIK3CA*, enrichment to 21% was observed, which was 210-fold higher enrichment than in the initial sample and was sufficient for the subsequent analyses. In contrast to the correlation of the enrichment efficiency with the initial SNV alleles for *KRAS* and *PIK3CA*, the enrichment of the deletion mutation of *EGFR* in the triplex sample achieved a nearly 100% mutation frequency for a wide range of initial mutant alleles. Together, the results indicate that the direction of the orthogonal activity of *Pf*Ago by corresponding gDNAs provides a simple single-tube reaction for multiplex enrichment. We also noted that there were some differences in the enrichment efficiency among different variants in the multiplex system. The performance could be improved through empirical optimization and bioinformatic analysis, as the thermodynamic parameters for the designed gDNA pairing target are under investigation in our lab.

## Discussion

In this study, we described a *Pf*Ago-directed PCR method for rapid and efficient enrichment and supersensitive detection of rare variant alleles. Our technique, which we called A-Star, significantly improves the sensitivity for rare variant alleles detection: This method enables specific amplification of SNV allele fractions as low as 0.01% with an over 5500-fold efficiency (Fig. 3a) and detection of rare SNV at a limit of single-digit copies of rare SNV as tested with the mocked cfDNA sample of 0.1% VAF (Fig. 3d). A major advantage of our approach is the combined ability of precise gDNA-target DNA pairing, discriminated DNA cleavage by *Pf*Ago endonuclease at DNA melting temperature, and variant alleles amplification in a single tube experiment (Fig. 1). Importantly, the greatly improved detection sensitivity relies on the basis of a very easy-to-implement design concept and is ideal for targeting virtually any mutant sequences. With programmable WT cleavage and variant alleles exponential amplification, A-Star can spare the use of RPA which loosely control the rate of amplification, less efficiency for amplifying highly structured regions, and challenges for primer design. Except derived from broadly used conventional PCR, the A-Star uses short DNA fragments as guides rather than RNAs in CRIPSR-system and thus provides a cheap and favorable way to synthesize and develop assay for variant alleles assay. Moreover, A-Star can provide multiplex availability with *Pf*Ago solely directed by the corresponding gDNAs pool in a single-tube reaction (Fig. 5), which substantially saves the labor and time for screening the multiple enzyme candidates used in Cas13a system and thus lowers the assay cost.

Oncogenes analysis have revealed that the cellular genome consists of pools of SNVs and indel mutations^32^, which are proved to affect the normal cell metabolism and signal transduction, such as *KRAS* mutation affects the Ras signaling pathway and holds the key clue in lung and colorectal carcinomas’ onset and development^33^. It is noteworthy that many patients’ ctDNA levels are considerably lower (<0.5%), whereas ctDNA concentrations typically drop following therapy^34, 35^. However, existing variant alleles detection techniques are still unsatisfactory in terms of their sensitivity, cost, time and simplicity, such as sanger sequencing and PCR-based methods are limited to their detection sensitivity and multiplex difficulty, while deep sequencing approaches are less economical for operation complexity and redundant read capacity on WT and non-targeted sequence. Our method improves the sensitivity of rare variant alleles detection (0.01% VAF), allowing the detection of many important variant alleles that are missed with current methods. Compared with qPCR and ARMS-PCR with approximately 20% detection limit, our method improves the detection sensitivity by two orders of magnitude, which favors the clinical detection for rare variant alleles for cancer diagnosis.

A-Star possesses an extra-high variant alleles enrichment efficiency since *Pf*Ago is compatible with PCR, thus combining the advantage of WT elimination with simultaneous variant alleles amplification in single tube, in contrast with the two or three round PCR enrichment methods of CUT-PCR (Cas9-based) and NAVIGATER (*Tt*Ago-based)^23, 36^. In the CUT-PCR and NAVIGATER methods, a procedure including PCR for DNA enrichment and target DNA cleavage was performed repeatedly for obtaining the high detection efficiency and sensitivity. However, only 20-fold and 60-fold enrichment was obtained after 3 rounds of CUT-PCR or 2 rounds of NAVIGATER, respectively^23, 36^. In this study, the single-tube A-Star was proved to achieve more efficient enrichment of rare variant alleles than the uncoupled cleavage and PCR procedure (Fig. 3 a and b, 75-fold vs 6-fold enrichment for 1% VAF sample, whereas 610-fold vs non enrichment for 0.1% VAF sample), demonstrating once again the strong superiority of the compatibility of the specific cleavage and routine amplification during PCR process.

Considering the possibility of multiple gene mutations existed in the ctDNA sample, such as *EGFR, KRAS*, or *PIK3CA, et al*, in lung carcinoma, the multiplex detection is necessary for guiding the use of targeted therapy by identifying precisely the mutated genotype. Here, A-Star confers advantages of providing multiplex enrichment and detection, with integration of multiple gDNAs, *Pf*Ago cleavage and PCR amplification in a single tube. We observed successfully multiplex variant alleles detection with *Pf*Ago’s orthogonal activity, that was proved by non-interfering signals observed using the duplex and triplex samples (Fig. 5b). We also demonstrated multiplex detection of A-Star for three oncogenes in the same reaction with the input of mocked cfDNA standard of 0.1% VAF (Fig. 5c). Together, these indicate that A-Star can be for use in simultaneously analyzing multiple instances and classes of somatic mutations in a single, low-cost and rapid reaction, notably, by adding multiple gDNA fragments and primers without further optimization. This is a marked difference from the CRISPR-based multiplex detection system SHERLOCK v2, which requires both the selection of multiple enzymes and the optimization of reaction parameters^24^. We also noted that there were some differences in the enrichment efficiency for the different rare variant alleles in the multiplex system that caused by rules governing the precise discrimination of the WT and variant alleles with varied sequences. This performance could be improved through empirical optimization and rational bioinformatic analysis, as the thermodynamics parameters of the designed gDNA pairing target are under investigation in our lab.

Although we here focus on applications related to oncogenes, our method can be applied for any situation that requires genotyping of rare alleles from complex samples, in which rare variants display disproportionate impact, such as infectious diseases caused by antibiotic-resistant subpopulations. Additionally, through multiplexing detection in a single reaction, we can foresee A-Star to facilitate much broader spectrum of basic research and vast clinical utility.

## Methods

### Nucleic Acid Preparation

The ssDNA target, gDNA, and primers were synthesized commercially (Sangon Biotech, China). The 626 bp dsDNA target was synthesized by GenScript (Nanjing, China) in the form of the pET-28a-derived plasmid. The plasmid DNA was extracted using a Plasmid DNA MiniPreps Kit (Generay, China). The 130-160 bp dsDNA target was obtained by amplification of the corresponding plasmid with the primers designed using NCBI Primer-BLAST, with parameters set for amplicon size between 130 and 160 nt, primer melting temperatures between 50 °C and 60 °C, and primer sizes between 18 and 25 nt. For PCR amplification, 2X PCR Precision™ MasterMix (Abm, Canada) was used. The target dsDNA fragments for the titration experiments were quantified using a PikoGreen dsDNA Quantitative Kit (Life iLab Biotech, China). All of the nucleic acids used in this study are listed in Table S1-S4.

### *Pf*Ago Expression and Purification

A codon-optimized version of the *Pf*Ago gene was synthesized by GenScript (Nanjing, China), which was designed as the pET28a-derived plasmid pEX-*Pf*Ago with an N-terminal His-tag. The expression plasmid was transformed into *E. coli* BL21(DE3) cells. A 5 mL seed culture was grown at 37 °C in LB medium with 50 μg/mL kanamycin and was subsequently transferred to 1 L of LB in a shaker flask containing 50 μg/mL kanamycin. The cultures were incubated at 37 °C until the OD_600_ reached 0.8-1.0, and protein expression was then induced by the addition of isopropyl β-D-thiogalactopyranoside (IPTG) to a final concentration of 1 mM, followed by incubation for 16 h at 20 °C. Cells were harvested by centrifugation for 20 min at 6,000 rpm, and the cell pellet was collected for later purification. Cell pellets were resuspended in lysis buffer (20 mM Tris/HCl, 1 M NaCl, pH 8.0) and then disrupted using a High Pressure Homogenizer at 600-800 bar for 3 min (Gefran, Italy). The lysates were centrifuged for 30 min at 12,000 rpm at 4 °C, after which the supernatants were subjected to Ni-NTA affinity purification with elution buffer (20 mM Tris/HCl, 1 M NaCl, 200 mM imidazole, pH 8.0). Further gel filtration purification using Superdex 200 (GE Tech, USA) was carried out with elution buffer (20 mM Tris/HCl, 1 M NaCl, pH 8.0). The fractions resulting from gel filtration were analyzed by SDS-PAGE, and fractions containing the protein were flash frozen at −80°C in storage buffer (20 mM Tris-HCl, pH 8.0, 250 mM NaCl, 10% (v/v) glycerol).

### DNA Cleavage Assays

Generally, *Pf*Ago-mediated cleavage assays were carried out in reaction buffer (15 mM Tris/HCl pH 8.0, 250 mM NaCl, and 0.5 mM MnCl_2_)^28^. For ssDNA cleavage, 0.2 µM *Pf*Ago, 2 µM gDNA, and 0.8 µM ssDNA target were mixed in reaction buffer and then incubated for 15 min at 95 °C in a thermocycler (Eppendorf, Germany). Following high-temperature incubation, the samples were cooled down by slowly lowering the temperature at a rate of 0.1 °C/s until it reached 10°C. The reactions were stopped via the addition of loading buffer (95% formamide, 0.5 mmol/L EDTA, 0.025% bromophenol blue, 0.025% xylene cyanol FF) at a 1:1 ratio (v/v); the samples were then separated in 16% denaturing polyacrylamide gels and analyzed by staining with GelRed (Biotium, USA). The nucleic acids were visualized using a G:BOX Chemi imager (Syngene, USA), and data were analyzed using Quantity One (Bio-Rad, USA). For the dsDNA cleavage assays, 0.16 µM *Pf*Ago, 2 μM gDNAs, and 58 nM dsDNA target were mixed in reaction buffer before incubation for 15 min at 95 °C in a thermocycler (Eppendorf, Germany). Following this high-temperature incubation step, the samples were cooled down by slowly lowering the temperature at a rate of 0.1 °C/s until it reached 10 °C. The reactions were quenched with 5× DNA loading buffer (Generay, China) and analyzed in 2% agarose gels. The gels were visualized with a Fuji FLA7000 scanner (FUJIFILM Life Science, USA), and the data were analyzed using Quantity One (Bio-Rad, USA).

### *Pf*Ago Activity at Different Temperatures

The effects of temperature on *Pf*Ago activity mediated by different guides were tested across a range of temperatures from 55 to 99 °C. Generally, 2.5 µM *Pf*Ago, 2 µM ssDNA guide, and 0.8 µM ssDNA target were mixed in reaction buffer, and the mixture was then incubated for 30 min at 54.8 °C, 58.3 °C, 60.7 °C, 66.0 °C, 71.0 °C, 74.9 °C, 79.9 °C, 85.1 °C, 90.2 °C, 95.1 °C, 96.9 °C, or 98.7 °C. The samples were resolved in 16% denaturing polyacrylamide gels. The gels were stained using GelRed (Biotium, USA). The nucleic acids were visualized using a G:BOX Chemi imager (Syngene, USA), and the data were analyzed using Quantity One (Bio-Rad, USA).

### Preparation of Variant Allele Fraction (VAF) Samples

Approximately 160 bp WT and mutant DNA fragments for each of *PIK3CA* E545K, *KRAS* G12D, and *EGFR* delE746-A750 were obtained by PCR amplification. Then, the PCR products were purified using a GeneJET Gel Extraction Kit (Thermo Scientific, USA) and quantified with PikoGreen dsDNA quantitative kits (Life ilab Bio, China). These wild-type and mutant DNA fragments were mixed to obtain VAFs of 1%, 0.1%, 0.01% for each SNV allele in a total DNA concentration of 10 nM for the evaluation of the enrichment performance of A-Star.

### Coupling cleavage and enrichment of A-Star and the uncoupling process

For the *Pf*Ago-coupled PCR process, 30 nM *Pf*Ago, 600 nM gDNA, 10 nM target DNA with 0.1% and 1% VAF of KRAS G12D, forward and reverse primer at 200 nM, and 500 µM Mn^2+^ were mixed in 2X PCR Taq MasterMix (Abm, Canada). Enrichment proceeded in a Mastercycler® RealPlex instrument (Eppendorf, Germany) with a temperature profile of 94 °C for 3 minutes, followed by 25 cycles of amplification (94 °C for 30 seconds, 55 °C for 30 seconds, and 72 °C for 20 seconds) and a final 72 °C extension for 1 minute. For the uncoupling process, the reaction mixture is the same with *Pf*Ago-coupled PCR process. Enrichment proceeded in the same instrument with a temperature profile of 94 °C for 15 minutes, then the cleavage products were cleaned up and followed by 25 cycles of amplification as procedure described above.

### A-Star for Variant Alleles Enrichment

For the *Pf*Ago-coupled PCR system, 30 nM *Pf*Ago, 600 nM gDNA, 10 nM target DNA with different VAFs, each of forward and reverse primer at 200 nM, and 500 µM Mn^2+^ were mixed in 2X PCR Taq MasterMix (Abm, Canada). Enrichment proceeded in a Mastercycler® RealPlex instrument (Eppendorf, Germany) with a temperature profile of 94 °C for 3 minutes, followed by 25 cycles of amplification (94 °C for 30 seconds, 55 °C for 30 seconds, and 72 °C for 20 seconds) and a final 72 °C extension for 1 minute. For variant alleles enrichment in mock cell free-DNA (cfDNA), mock cfDNA standards simulating actual patient cfDNA samples were purchased from a commercial vendor (Horizon Discovery Group, UK). These standards were provided for each target (*PIK3CA* E545K, *KRAS* G12D, and *EGFR* del) with different VAFs (0.1%, 1%, and 5%) at a concentration of 33.3 ng/µl. Then, 1 µl of these standards was used as input to perform preamplification and subsequently examined using A-Star. The preamplification procedure was carried out in a 25 µl reaction volume using 33.3 ng of mock cfDNA, 2X PCR Precision™ MasterMix (Abm, Canada), and each forward and reverse primer at 250 nM, using a Mastercycler® RealPlex instrument (Eppendorf, Germany) with a temperature profile of 94 °C for 3 minutes, followed by 30 cycles of amplification (94 °C for 10 seconds, 55 °C for 30 seconds, and 72 °C for 20 seconds) and a final 72 °C, extension for 1 minute. Two microliters of the preamplification products was used as input to perform A-Star. The A-Star system consisted of 20-30 nM *Pf*Ago, gDNAs at a concentration 20-fold that of *Pf*Ago, each forward and reverse primer at 200 nM, and 500 µM Mn^2+^, which were mixed in 2X PCR Taq MasterMix (Abm, Canada). Enrichment proceeded in a Mastercycler® RealPlex instrument (Eppendorf, Germany) as described above. For variant alleles enrichment in tissue samples, 33 ng of extracted genomic DNA from each tissue sample was used as the input to perform preamplification, and the product was then examined using A-Star as described above for the variant alleles enrichment of mock cfDNA. For variant alleles enrichment in cfDNA samples, due to the low concentration of extracted cfDNA, 3.3 ng of extracted genomic DNA from each sample was used as the input to perform preamplification, and then 2 µl of preamplification products was used as the input to perform A-Star. The A-Star system consisted of 10-20 nM *Pf*Ago, gDNAs at a concentration 20-fold that of *Pf*Ago, each forward and reverse primer at 200 nM, and 500 µM Mn^2+^, which were mixed in 2X PCR Taq MasterMix (Abm, Canada). Enrichment proceeded in a Mastercycler® RealPlex instrument (Eppendorf, Germany) as described above.

### Triplex Variant Alleles Enrichment of A-Star

Triplex enrichment was performed in 25 µL reaction volumes with 200 nM *Pf*Ago, each gDNA at 4000 nM, 500 µM Mn^2+^, each forward and reverse primer at 200 nM, 0.4 mM dNTPs (Sangon, China), 0.5 μl of Taq DNA Polymerase (Abm, Canada), and three VAF targets each at a concentration of 10 nM in 10X PCR buffer (Abm, Canada). The reactions were carried out in a Mastercycler® RealPlex instrument (Eppendorf, Germany) with a temperature profile of 94 °C for 5 minutes, followed by 25 cycles of amplification (94 °C for 30 seconds, 55 °C for 30 seconds, 72 °C for 20 seconds) and a final 72 °C extension for 1 minute. For triplex SNV enrichment of mock cfDNA, cfDNA standards purchased from a commercial vendor (Horizon Discovery Group, UK) were used as the three targets (*PIK3CA* E545K, *KRAS* G12D, and *EGFR* del) with different VAFs (0.1%, 1%, and 5%) at a concentration of 33.3 ng/µl. Then, 1 µl of these standards was used as the input to perform preamplification, followed by analysis using A-Star. The preamplification process was carried out in a 25 µl reaction volume using 33.3 ng of mock cfDNA, 2X PCR Precision™ MasterMix (Abm, Canada), and each forward and reverse primer at 250 nM for three targets, using a Mastercycler® RealPlex instrument (Eppendorf, Germany) with a temperature profile of 94 °C for 3 minutes, followed by 30 cycles of amplification (94 °C for 10 seconds, 55 °C for 30 seconds, and 72 °C for 20 seconds), and a final 72 °C, extension for 1 minute. Then, 2 μl of the preamplification product was used as the input to perform A-Star as described above for the SNV enrichment of mock cfDNA.

### Detection of Enriched Products by Sanger Sequencing and TaqMan Real-Time PCR

The A-Star enriched products were checked for quality and yield by running 5 µl of the products in 2.0% agarose gels and visualized on a Fuji FLA7000 scanner (FUJIFILM Life Science, USA), then processed directly for Sanger sequencing (Sangon, China). The primers and probes for the targets (*PIK3CA* E545K, *KRAS* G12D, and *EGFR* del) were designed using Beacon Designer’s standard assay design pipeline and ordered as individual primers and probes from Life Technologies (Thermo Fisher, USA). To validate the enriched mutant products, assays were performed using AceQ® qPCR Probe Master Mix (Vazyme biotech co., Ltd), and the results were quantified in a StepOnePlus™ Real-Time PCR System (Thermo Fisher) with the primer and probes sets (Supplementary Data Table S4) at final concentrations of 250 nM primers and 200 nM probes. The temperature profile for amplification consisted of an activation step at 95 °C for 8 minutes, followed by 40 cycles of amplification (95 °C for 15 seconds and 60 °C for 45 seconds).

### Collection and DNA extraction of patient samples

Patients with cancers of the pancreas, colorectum, lung, or breast were recruited from Shanghai General Hospital under methods approved by the Human Research Committee of Shanghai General Hospital. The ‘healthy donor’ samples consisted of peripheral blood samples obtained from 2 individuals with no history of cancer. The cancer and healthy control samples listed in Table S5 were processed in an identical manner. DNA extraction from formalin-fixed, paraffin-embedded (FFPE) tumor blocks was performed using a QIAamp® DNA Blood Mini Kit (Qiagen, Germany) according to the recommended kit protocol. Five milliliters of peripheral venous blood were collected in Streck Cell-Free DNA BCT (Streck Inc., USA), followed by centrifugation at 1,600 g for 10 min at 10 °C, and the resulting supernatant was clarified by additional centrifugation. Clarified plasma was transferred to a fresh tube, and DNA was immediately extracted using a QIAamp® DNA Blood Mini Kit (Qiagen, Germany). Finally, exome sequencing and data processing to produce a BAM file were performed using established NGS analytical pipelines at Shanghai General Hospital to validate the SNV alleles of the samples.

### Reporting summary

Further information on research design is available in the Nature Research Reporting Summary linked to this article.

## Data availability

The source data underlying Figs. are provided as a Sources Data file. All other data are available from the corresponding author (Y. F.) upon reasonable request.

## Acknowledgements

We thank Ming Yu from Shanghai Jiao Tong University for helpful comments on the manuscript. This research was supported by the grants from the Natural Science Foundation of China (31770078) and Ministry of Science and Technology (2017YFE0103300).

## Author contributions

Y.F. and Q.L. conceived the project and designed the experiments. Q.L., X.G., G.H.X., and Y.S.C. carried out the biochemical experiments. Z.L.L. designed the TaqMan specific probes and prepared the clinical samples. L.T.Y, K.L. and H.X.W. assisted with TaqMan analysis, system optimization and clinical experiment, respectively. Q.L., S.K.L, K.L., Z.X.D. and Y.F. wrote the manuscript with contributions from all authors. All authors read and commented on the manuscript.

## Additional information

### Supplementary Information

accompanies this paper at:

