## Supplementary Figs for "Argonaute-mediated system for supersensitive and multiplexed detection of rare mutations"

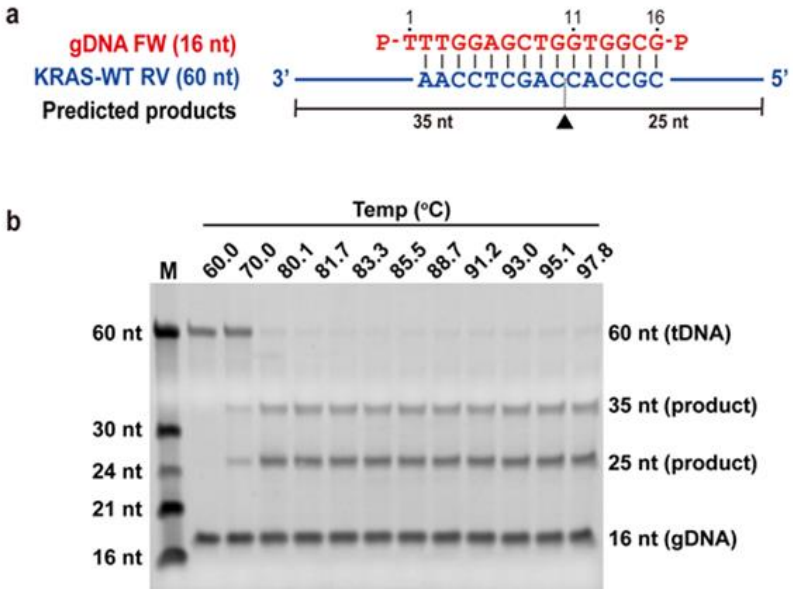


**Figure S1. Evaluation of the temperature dependence of *Pf*Ago cleavage activity on the ssDNA target.** (**a**) The sequences of the ssDNA target and gDNA. (**b**) Gel electrophoresis of cleavage products from the ssDNA target. *Pf*Ago was incubated with a 16 nt gDNA and 60 nt ssDNA of *KRAS* G12D as a target at a 1:10:1 molar ratio (*Pf*Ago:gDNA:tDNA) at various temperatures for 15 min with 0.5 mM $Mn^{2+}$. The cleavage products were resolved in denaturing polyacrylamide gels. M: ssDNA Marker; nt: nucleotides.

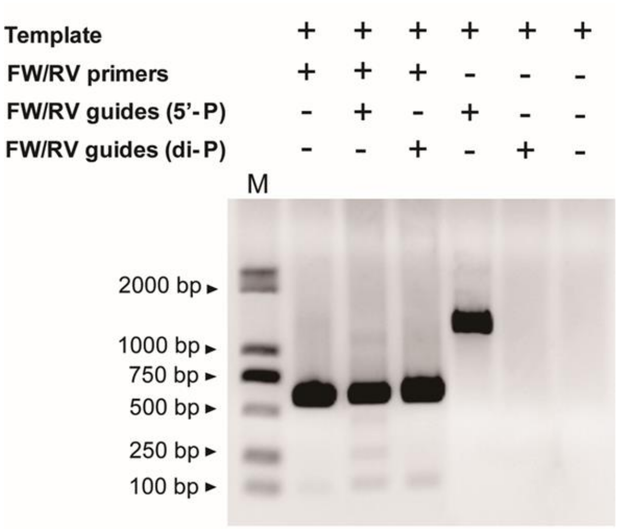


**Figure S2. Evaluation of the di-phosphate gDNA dependence of *Pf*Ago cleavage activity on the ssDNA target.** A plasmid containing *KRAS* G12D as the template was used for PCR. A pair of primers was designed for the 626 bp of *KRAS* G12D amplicon, and a pair of gDNAs were designed with either 5’-P or 5’, 3’-di-P modifications across the *KRAS* G12D SNV site. Thermal cycling started with a preincubation step at 94 °C for 3 mins for polymerase activation, followed by 30 PCR cycles (94 °C for 30 s, 55 °C for 30 s, and 72 °C for 20 s). The products were resolved in an agarose gel. M: dsDNA Marker; bp: base pairs; 5’-P: only the 5’- termini is modified by phosphate group; di-P: both the 5’- and 3’- termini are modified by phosphate groups.

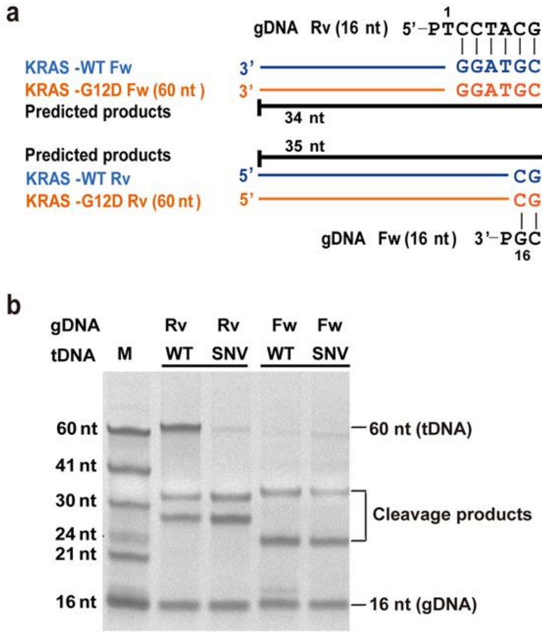


**Figure S3. Comparison of the cleavage efficiency of WT and SNV ssDNA targets as directed by gDNA complementary to the WT sequence.** (**a**) Schematic diagram of the gDNAs and the ssDNA targets in both the forward and reverse strands of *KRAS* G12D WT and SNV. (**b**) Gel electrophoresis for the cleavage products of both strands of *KRAS* G12D WT and SNV as directed by the corresponding gDNAs. *Pf*Ago was incubated with 16 nt gDNAs and 60 nt of WT or variant ssDNAs target at a 1:10:1 molar ratio (*Pf*Ago: gDNA: tDNA) at 95 °C for 15 min. The cleavage products were resolved in denaturing polyacrylamide gels. M: ssDNA marker; nt: nucleotides.

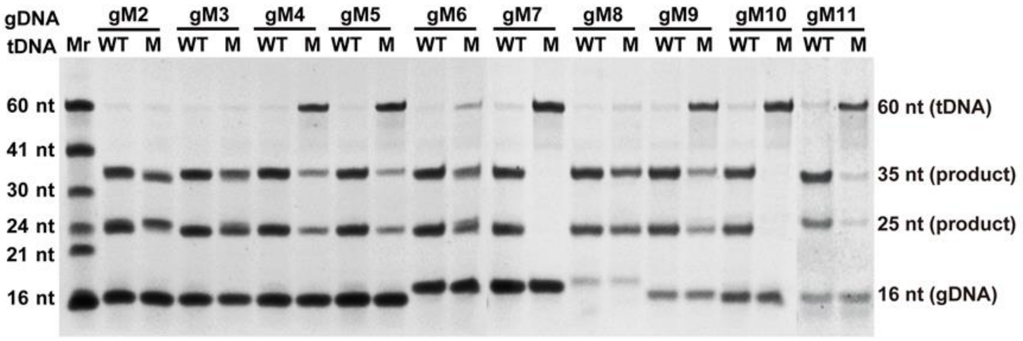


**Figure S4. Discriminating cleavage of the SNV ssDNA target using gDNAs with an extra mismatch.** Electropherogram of the cleavage products of the *KRAS* G12D WT and SNV ssDNA targets using the designed gDNAs. *Pf*Ago was incubated with 16 nt serial gDNAs containing mismatches from positions 2 to 11 and 60 nt of the WT and variant ssDNAs as targets at a 1:10:1 molar ratio (*Pf*Ago: gDNA: tDNA) at 95 °C for 15 min. The cleavage products were resolved in denaturing polyacrylamide gels. Mr: ssDNA marker; nt: nucleotides.

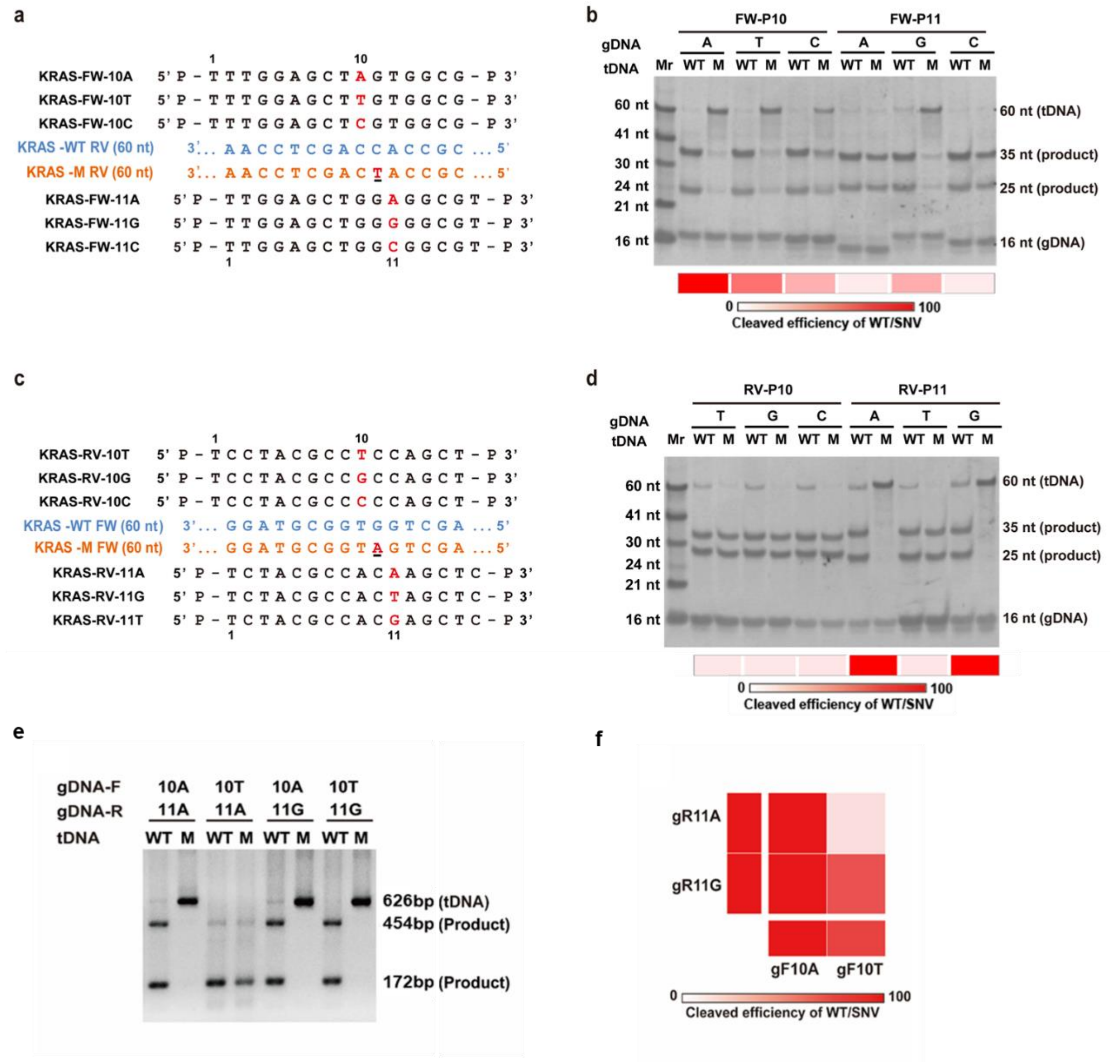


**Figure S5. Characterization of the discriminating cleavage of *KRAS* G12D targets directed by synthetic gDNA with nucleotide identity at the 10 or 11 position.** (**a**) and (**c**) show the design of the gDNAs with different nucleotide substitutions at position 10 or 11 for both strands of the ssDNA targets. Mismatches are shown in red with the positions indicated. (**b**) and (**d**) show the results of the gel electrophoresis of cleavage products for each strand of the ssDNA targets. (**e**) and (**f**) show the cleavage products for dsDNA targets and the corresponding heatmap of discriminating cleavage activity. *Pf*Ago was incubated with 16 nt gDNAs and the WT and SNV ssDNA or dsDNA targets in a 1:10:1 molar ratio (*Pf*Ago: gDNA: tDNA) at 95 °C for 15 min. The cleavage products were resolved in denaturing polyacrylamide gels (**b, d**) and agarose gels (**e**). The specificity index in the heatmap is calculated as the ratio of the cleavage efficiency of WT to that of SNV. Mr: ssDNA marker; M: SNV ssDNA or dsDNA target; nt: nucleotides; bp: base pairs.

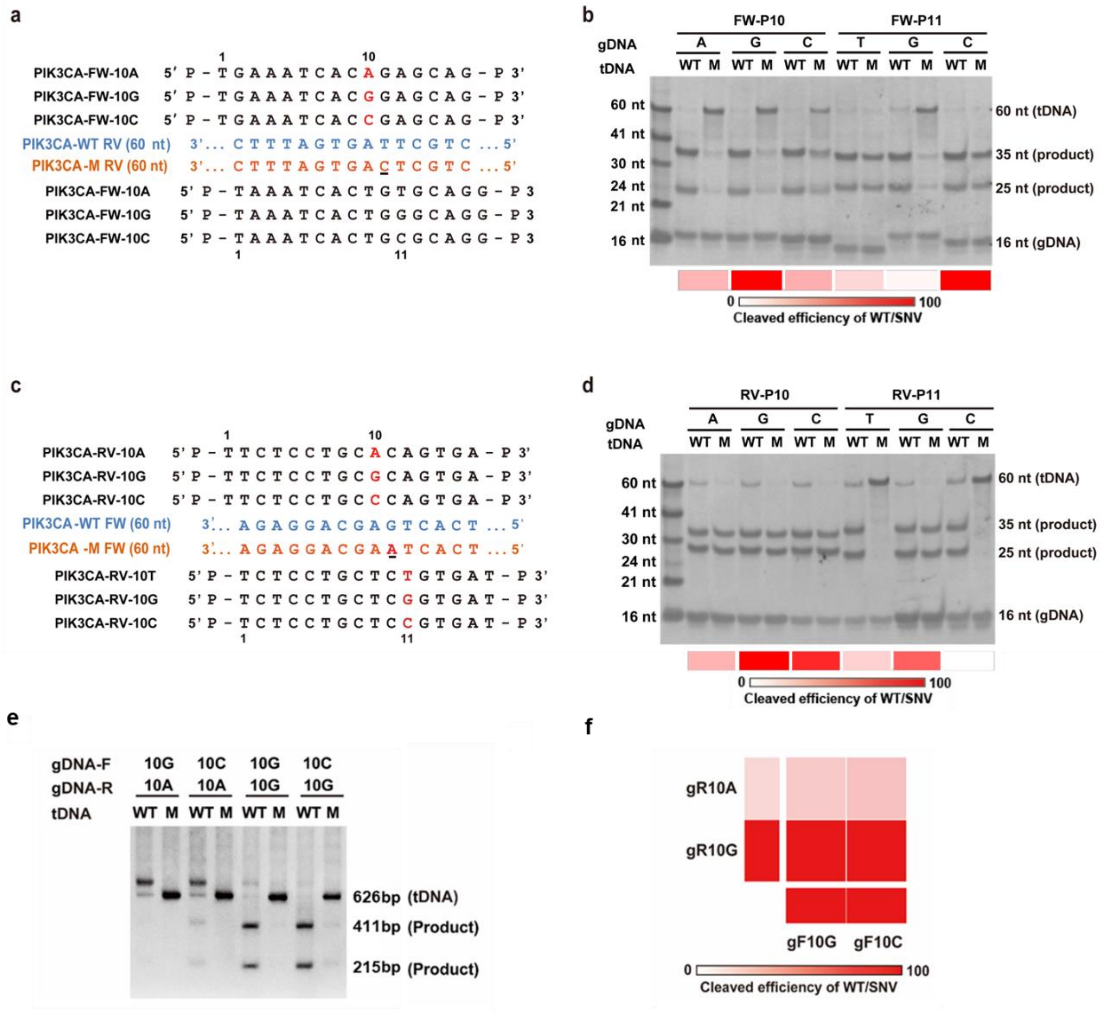

**Figure S6. Characterization of the discriminating cleavage of *PIK3CA* E545K targets directed by synthetic gDNA with nucleotide identity at the 10 or 11 position.** (**a**) and (**c**) show the design of gDNAs with different nucleotide substitutions at position 10 or 11 for both strands of the ssDNA targets. Mismatches are shown in red with the positions indicated. (**b**) and (**d**) show the results of gel electrophoresis of the cleavage products for each strand of the ssDNA targets. (**e**) and (**f**) show the cleavage products for the dsDNA targets and the corresponding heatmap of discriminating cleavage activity. *Pf*Ago was incubated with 16 nt gDNAs and the WT and SNV ssDNA or dsDNS targets at a 1:10:1 molar ratio (*Pf*Ago: gDNA: tDNA) at 95 °C for 15 min. The cleavage products were resolved in denaturing polyacrylamide gels (**b, d**) and agarose gels (**e**). The specificity index in the heatmap is calculated as the ratio of the cleavage efficiency of WT to that of SNV. Mr: ssDNA marker; M: SNV ssDNA or dsDNA target; nt: nucleotides; bp: base pairs.

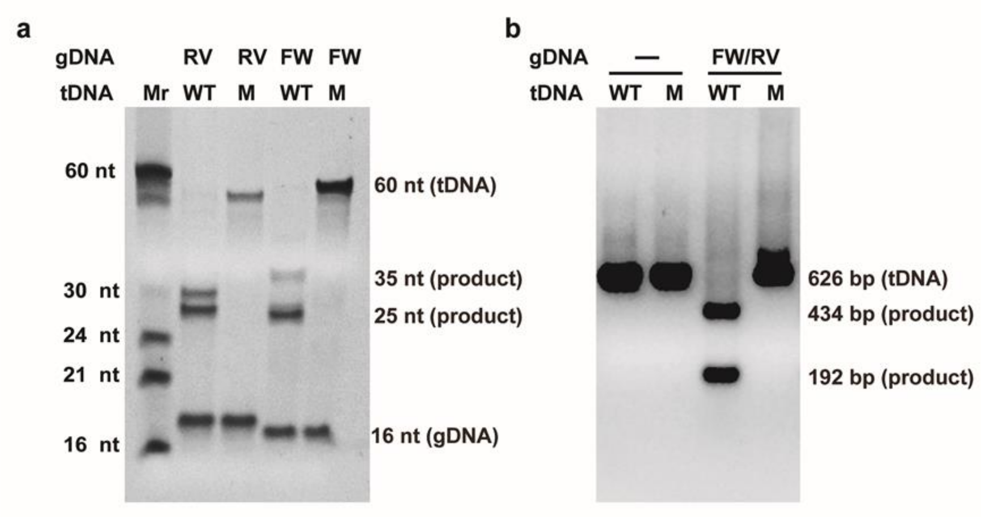


**Figure S7. Cleavage discrimination of an *EGFR* del target directed by gDNA completely matching the WT sequence.** (**a**) Cleavage efficiency for an *EGFR* del ssDNA target with a gDNA complementary to *EGFR* WT.(**b**) Cleavage efficiency for an *EGFR* del dsDNA target with a pair of gDNAs that completely paired with each strand of the WT dsDNA. *Pf*Ago loaded with the 16 nt gDNAs was incubated with the corresponding DNA target in a 1:10:1 molar ratio (*Pf*Ago: gDNA: tDNA) at 95 °C for 15 min. Nucleic acids were resolved in denaturing polyacrylamide gels (**a**) and agarose gels (**b**). Mr: ssDNA marker; M: mutant ssDNA or dsDNA target; nt: nucleotides; bp: base pairs.

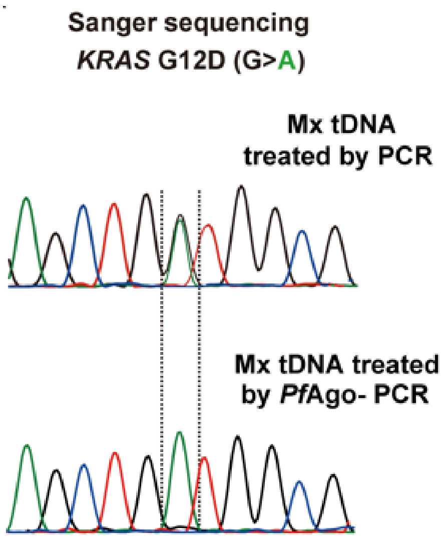


**Figure S8. Evaluation of the A-Star enrichment results for *KRAS* G12D through either PCR or *Pf*Ago-coupled PCR by Sanger sequencing.** A sample mixture composed of the *KRAS* WT and G12D SNV fragments at equal molar ratios was used as the input for either PCR or *Pf*Ago-coupled PCR. *Pf*Ago was incubated with the corresponding pair of gDNA targets at a 1:10 molar ratio (*Pf*Ago: gDNA). Thermal cycling started with a preincubation step at 94 °C for 3 mins for polymerase activation, followed by 30 PCR cycles (94 °C for 30 s, 55 °C for 30 s, and 72 °C for 20 s).

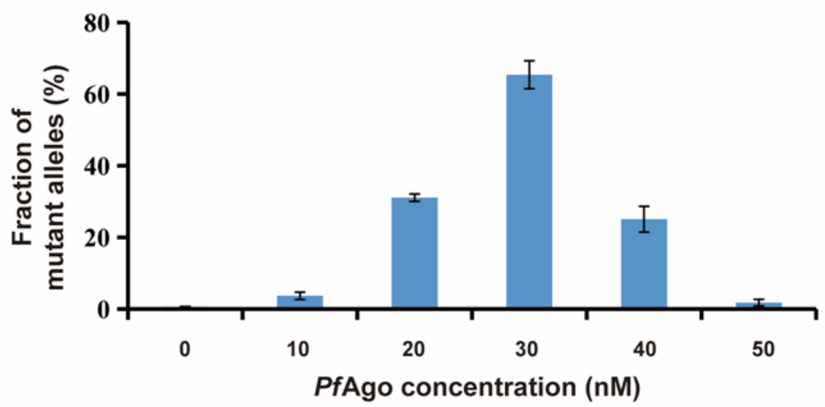


**Figure S9. Effect of the *Pf*Ago concentration on A-Star enrichment for a 1% VAF of *KRAS* G12D.** The *KRAS* G12D dsDNA target at a 1% VAF and 10 nM was tested with different *Pf*Ago concentrations but a consistent ratio of *Pf*Ago:gDNA (1:10) in the reactions. The enriched products were quantitatively measured using TaqMan real-time PCR. Error bars represent the mean ± s.d., n = 3.

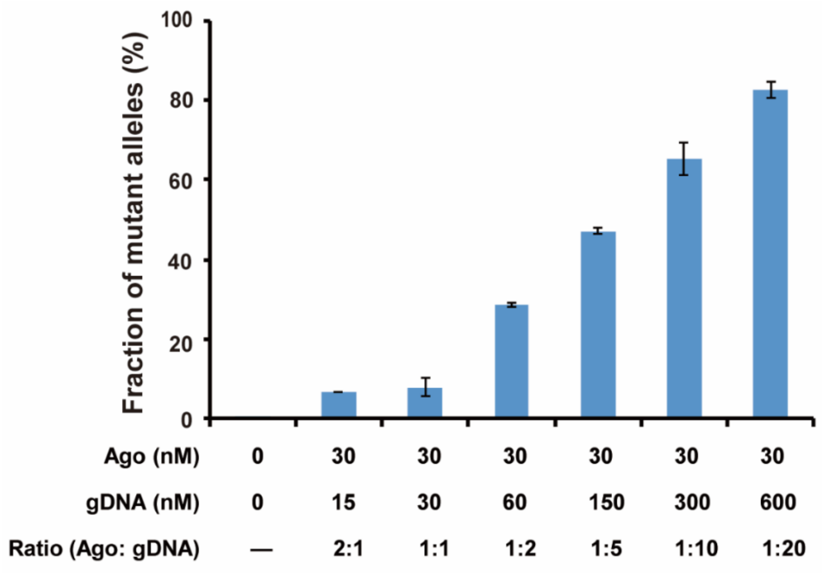


**Figure S10. Effect of the ratio of *Pf*Ago:gDNA on A-Star enrichment for a 1% VAF of *KRAS* G12D.** The dsDNA target with a 1% VAF of *KRAS* G12D at 10 nM was tested with different ratios of *Pf*Ago:gDNA but with a consistent 30 nM *Pf*Ago concentration in the reactions. The enriched samples were quantitatively measured using TaqMan real-time PCR. Error bars represent the mean ± s.d., n = 3.

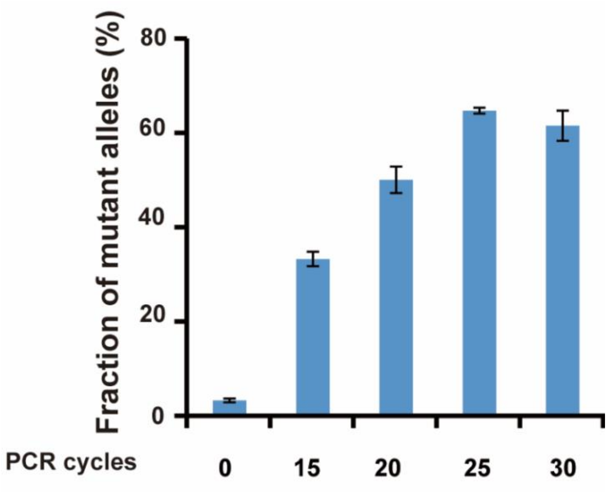


**Figure S11. Effect of the PCR program on A-Star enrichment for a 1% VAF of *KRAS* G12D.** The dsDNA target with a 1% VAF of *KRAS* G12D at 10 nM was tested using 30 nM *Pf*Ago and the *Pf*Ago:gDNA molar ratio was set to 1:20 in the reactions. Thermal cycling started with a preincubation step at 94 °C for 5 mins for polymerase activation and *Pf*Ago cleavage, followed by 15~30 PCR cycles (94 °C for 30 s, 55 °C for 30 s, and 72 °C for 20 s). The enriched products were quantitatively measured using TaqMan real-time PCR. Error bars represent the mean ± s.d., n = 3.

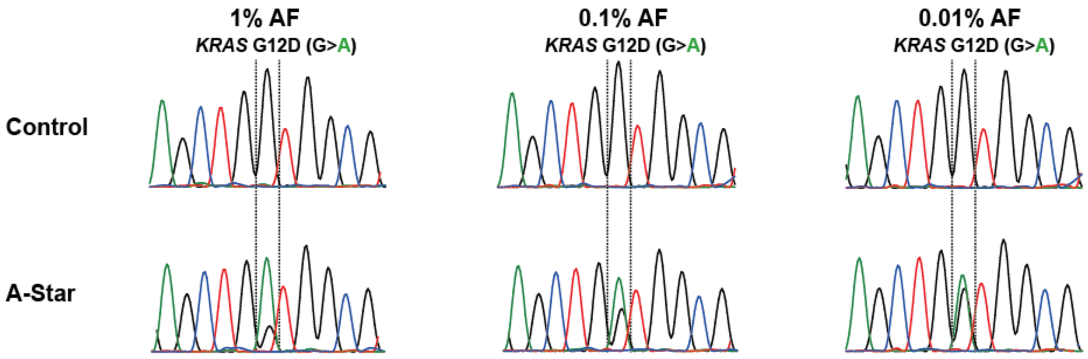


**Figure S12 Evaluation of the A-Star enrichment results for *KRAS* G12D with VAFs of 1%, 0.1%, and 0.01%.** The dsDNA target with different VAFs (1%, 0.1%, and 0.01%) of *KRAS* G12D at 10 nM was tested via the optimized A-Star treatment approach and then analyzed via Sanger sequencing. All the controls were preprocessed in the absence of a pair of gDNAs.

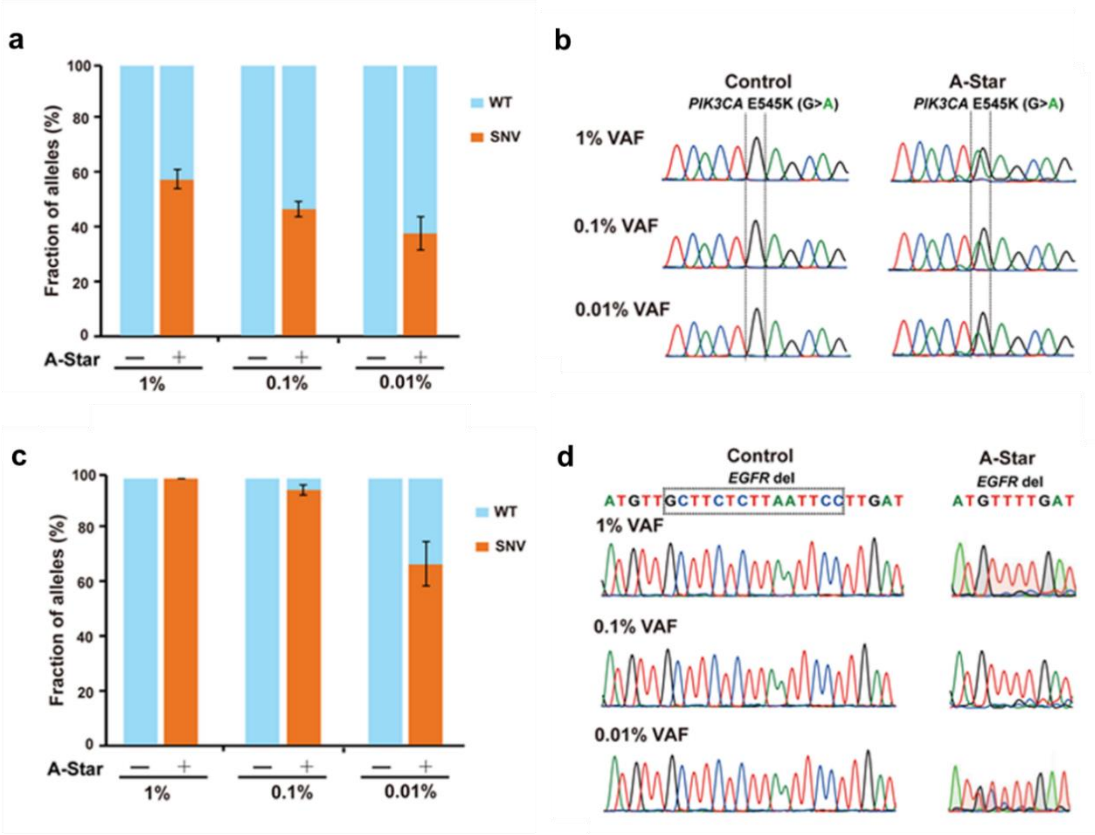


**Figure S13. Evaluation of the A-Star enrichment results for *PIK3CA* E545K and *EGFR* del with 1%, 0.1%, and 0.01% VAFs.** The synthetic target was originally mixed with the WT and mutant fragments at different VAFs at a 10 nM concentration. The samples were then used as the input for A-Star, followed by analysis using TaqMan real-time PCR and Sanger sequencing. TaqMan detection showed that A-Star could enrich rare variant alleles of *PIK3CA* E545K **(a)** and *EGFR* del **(c)** with a 0.01% VAF sensitivity. Error bars represent the mean ± s.d., n = 3. Strong signals were also observed for rare variant alleles of *PIK3CA* E545K **(b)** and *EGFR* del **(d)** according to the Sanger sequencing results. All the controls were preprocessed in the absence of a pair of gDNAs.

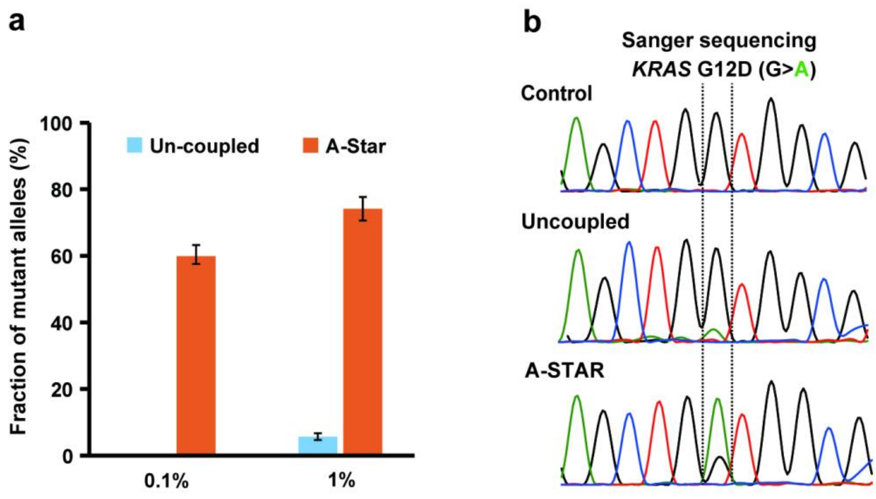


**Figure S14. Evaluation of the A-Star enrichment results for *KRAS* G12D via either the uncoupled or A-Star approach. (a)** Comparison enrichment efficiency of A-Star with the uncoupled *Pf*Ago-PCR tested with a 0.1% and 1% VAF sample of KRAS G12D by TaqMan PCR. **(b)** The dsDNA target with a 1% VAF of *KRAS* G12D at 10 nM was tested via the optimized A-Star treatment approach and then analyzed via Sanger sequencing. The controls were preprocessed in the absence of a pair of gDNAs.

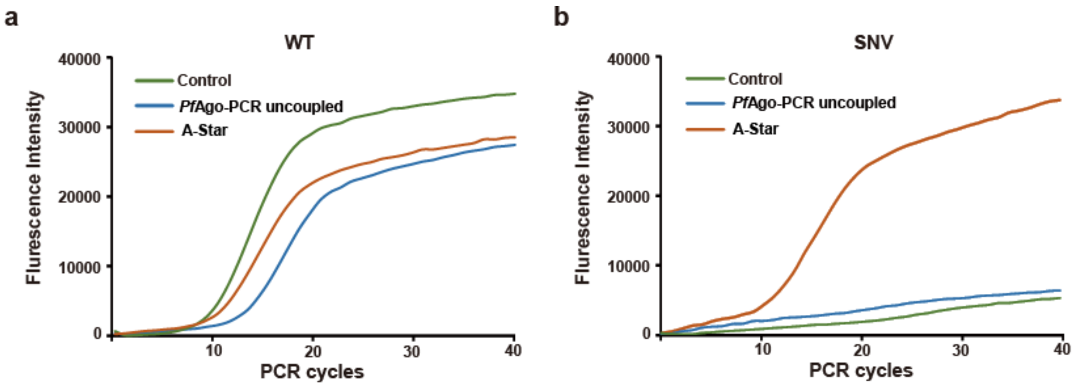


**Figure S15. Comparison of enrichment results for A-Star and the uncoupled *Pf*Ago-PCR reaction for a 1% VAF of *KRAS* G12D.** TaqMan real-time PCR amplification curves with a specific probe for the WT (**a**) and an SNV (**b**) targets. The dsDNA target with a 1% VAF of *KRAS* G12D at 10 nM was used for analysis with either optimized A-Star or the uncoupled *Pf*Ago-PCR reaction. The enriched samples were quantitatively measured using TaqMan real-time PCR. All the controls were preprocessed in the absence of a pair of gDNAs.

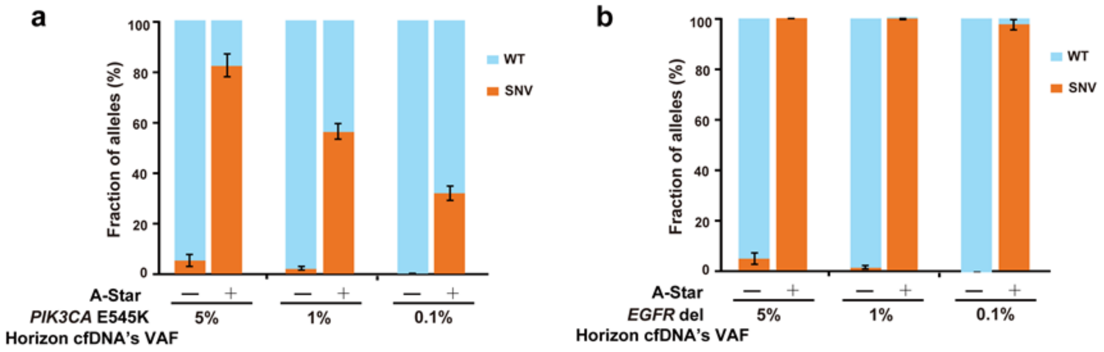


**Figure S16. Evaluation of the A-Star enrichment results for *PIK3CA* E545K (a) and *EGFR* del (b) Horizon cfDNA standard samples with different VAFs.** The standards were purchased from a commercial vendor (Horizon Discovery Group) with different VAFs of 0.1%, 1%, and 5% for *KRAS* G12D, *PIK3CA* E545K and *EGFR* del, respectively. 33 ng/μl of these standards was provided as the input to A-Star, followed by analysis using TaqMan real-time PCR. All the controls were preprocessed in the absence of a pair of gDNAs. Error bars represent the mean ± s.d., n = 3.

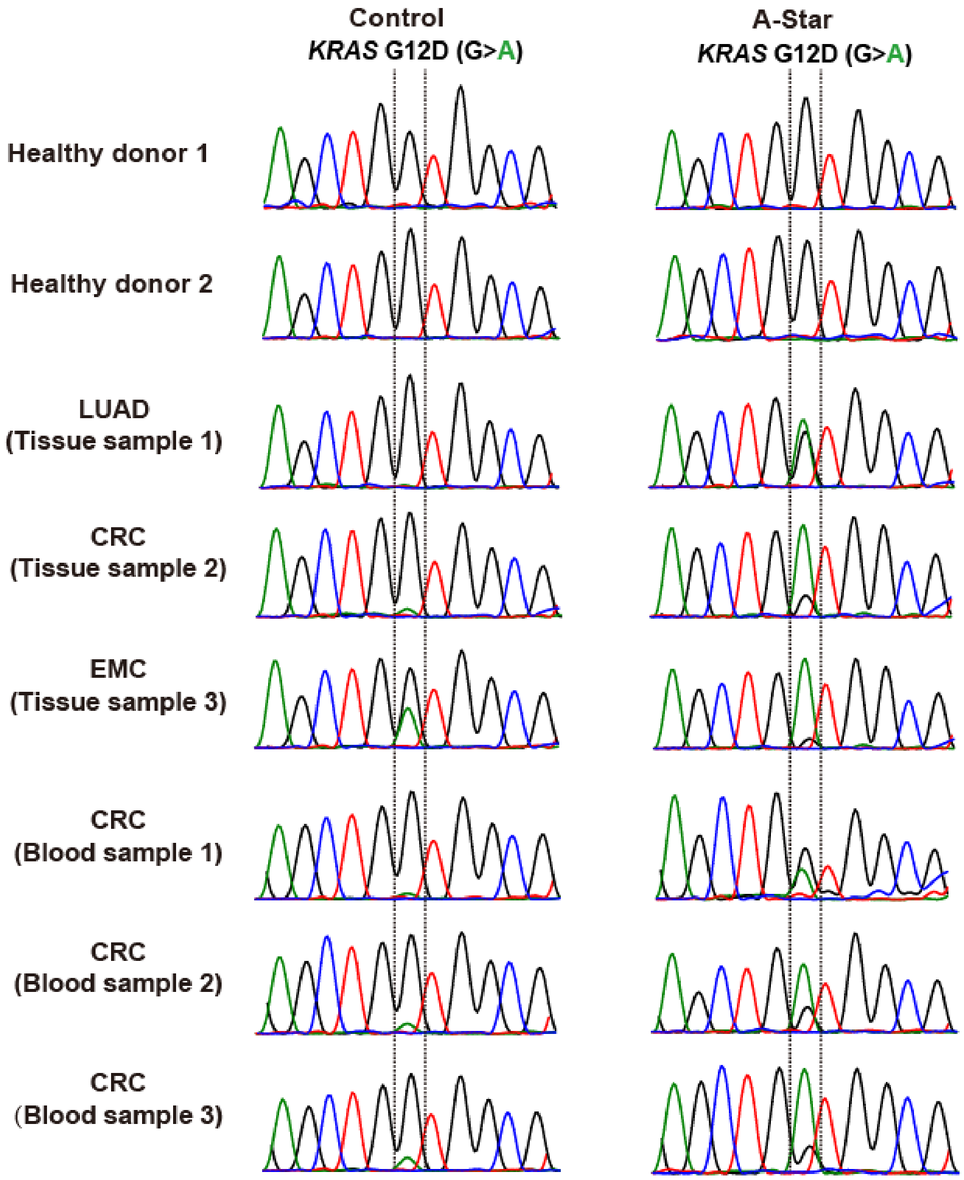


**Figure S17. Evaluation of the A-Star enrichment results for clinical samples of diverse cancer types by Sanger sequencing.** The clinical sample DNA was extracted and used as the input for A-Star, followed by analysis using Sanger sequencing. All the controls were preprocessed in the absence of a pair of gDNAs.

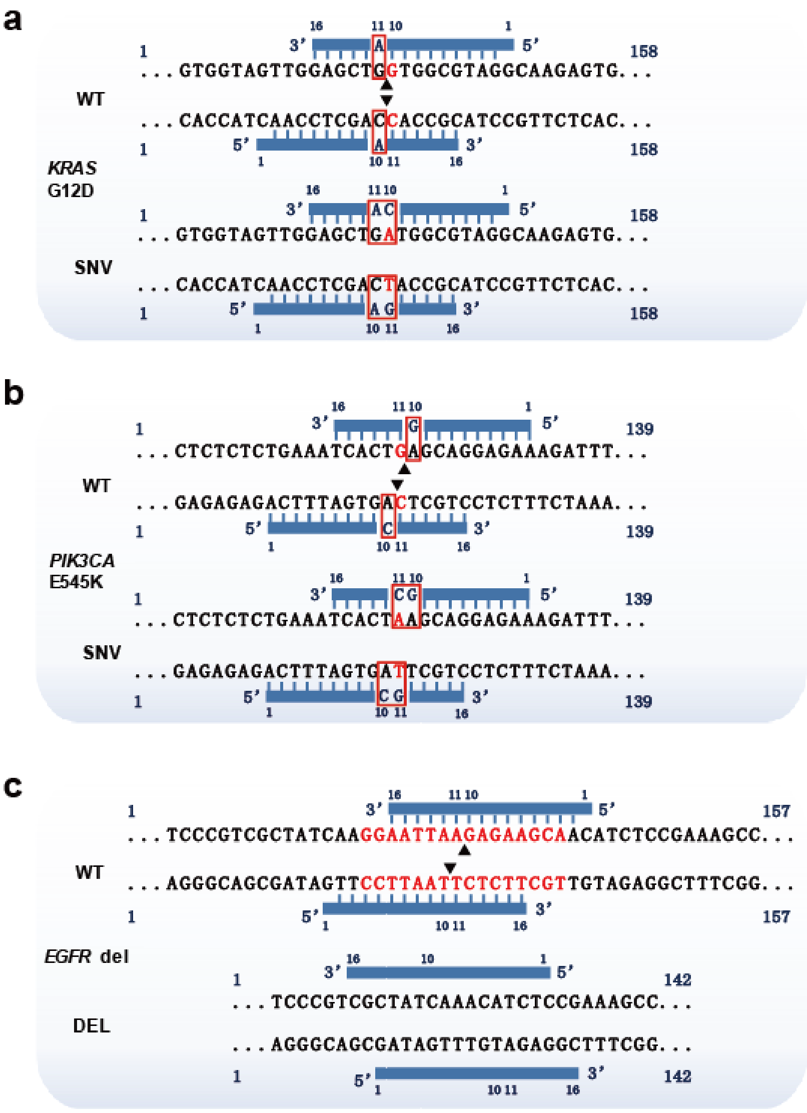


**Figure S18. Diagram of the target region and gDNA design for the multiplex detection of *KRAS* G12D, *PIK3CA* E545K, and *EGFR* del.** Mutation in the target are indicated in red; the mismatches introduced into the gDNAs are shown in red boxes; and the nucleotides deleted in *EGFR* del are highlighted in red.

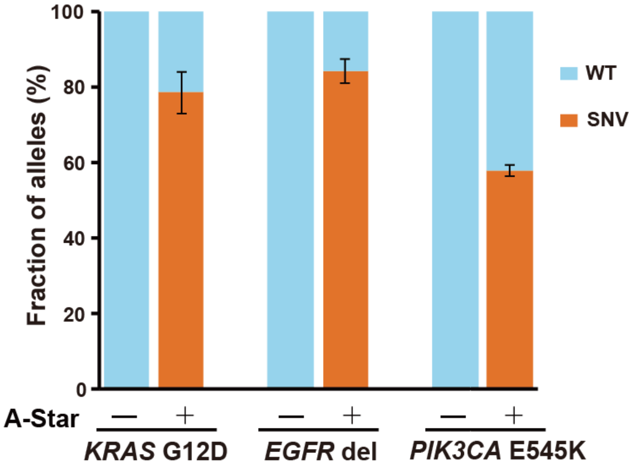


**Figure S19. Evaluation of the A-Star multiplexed enrichment results for *KRAS* G12D, *PIK3CA* E545K, and *EGFR* del with the 1% VAF sample.** The synthetic sample was mixed with a 1% VAF of *KRAS* G12D, *PIK3CA* E545K, and *EGFR* del at a 10 nM concentration and then used as the input for A-Star, followed by analysis with a TaqMan probe. All the controls were preprocessed in the absence of a pair of gDNAs. Error bars represent the mean ± s.d., n = 3.

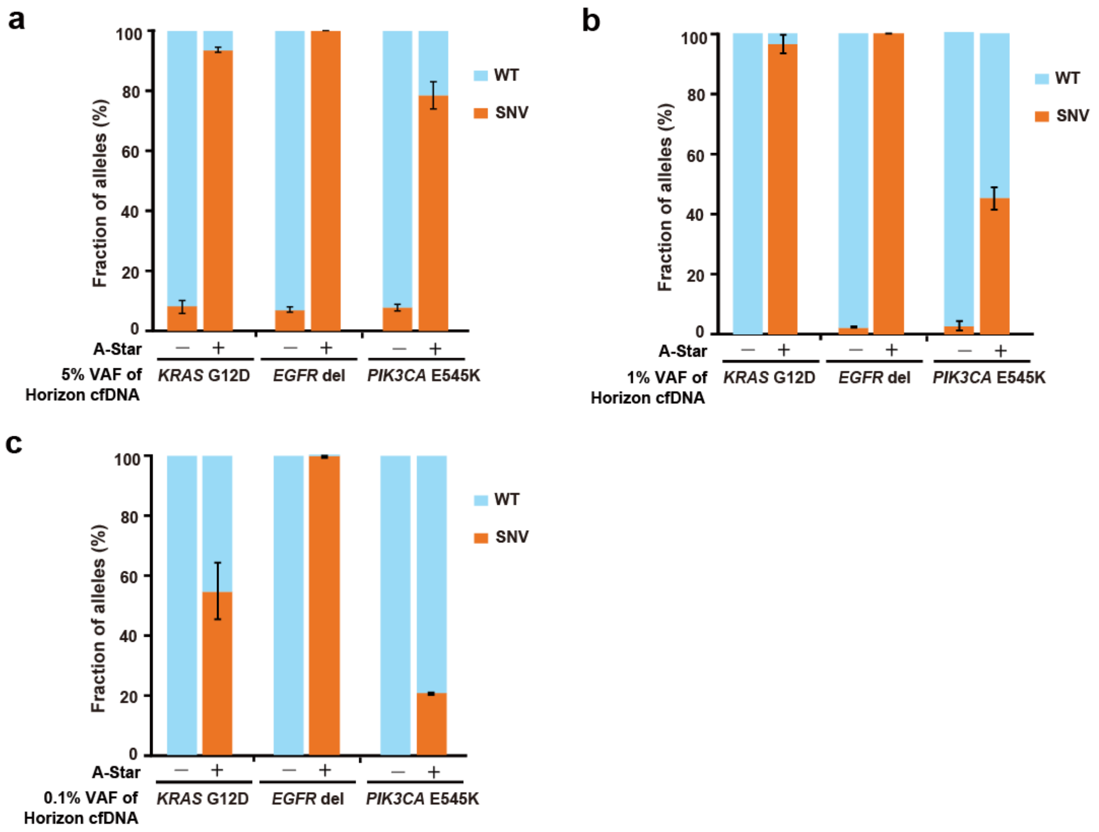


**Figure S20. Evaluation of the A-Star triplex enrichment results for *KRAS* G12D, *PIK3CA* E545K, and *EGFR* del Horizon cfDNA standard samples with different VAFs of 5% (a), 1% (b), and 0.1% (c).** The standards were purchased from a commercial vendor (Horizon Discovery Group) and had different VAFs of 0.1%, 1%, and 5% for *KRAS* G12D, *PIK3CA* E545K, and *EGFR* del, respectively. 33 ng/µl of these standards was used as the input to A-Star, followed by analysis via TaqMan real-time PCR. All the controls were preprocessed in the absence of pairs of gDNAs. Error bars represent the mean ± s.d., n = 3.

**Table S1. gDNA used in this study.**

| Name | gDNA sequence (5'→3') | Target | Presence |
|---|---|---|---|
| KRAS- FW | P-TTTGGAGCTGGTGGCG | KRAS-(WT)-RV/KRAS-(M)-RV | Fig. S1/Fig. S3 |
| KRAS- RV | P-TCCTACGCCACCAGCT | KRAS-(WT)-WT/KRAS-(M)-WT | Fig. S3 |
| KRAS_gM2 | P-TATGGAGCTGGTGGCG-P | KRAS-(WT)-RV/KRAS-(M)-RV | Fig. 2/Fig. S4 |
| KRAS_gM3 | P-TTAGGAGCTGGTGGCG-P | KRAS-(WT)-RV/KRAS-(M)-RV | Fig. 2/Fig. S4 |
| KRAS_gM4 | P-TTTCGAGCTGGTGGCG-P | KRAS-(WT)-RV/KRAS-(M)-RV | Fig. 2/Fig. S4 |
| KRAS_gM5 | P-TTTGCAGCTGGTGGCG-P | KRAS-(WT)-RV/KRAS-(M)-RV | Fig. 2/Fig. S4 |
| KRAS_gM6 | P-TTTGGTGCTGGTGGCG-P | KRAS-(WT)-RV/KRAS-(M)-RV | Fig. 2/Fig. S4 |
| KRAS_gM7 | P-TTTGGACCTGGTGGCG-P | KRAS-(WT)-RV/KRAS-(M)-RV | Fig. 2/Fig. S4 |
| KRAS_gM8 | P-TTTGGAGGTGGTGGCG-P | KRAS-(WT)-RV/KRAS-(M)-RV | Fig. 2/Fig. S4 |
| KRAS_gM9 | P-TTTGGAGCAGGTGGCG-P | KRAS-(WT)-RV/KRAS-(M)-RV | Fig. 2/Fig. S4 |
| KRAS_gM10 | P-TTTGGAGCTAGTGGCG-P | KRAS-(WT)-RV/KRAS-(M)-RV | Fig. 2/Fig. S4 |
| KRAS_gM11 | P-TCTACGCCACGAGCTC-P | KRAS-(WT)-FW/KRAS-(M)-FW | Fig. 2/ Fig. S4 |
| KRAS-RV-10T | P-TCCTACGCCTCCAGCT-P | KRAS-(WT)-FW/KRAS-(M)-FW | Fig. S4-S5 |
| KRAS-RV-10G | P-TCCTACGCCGCCAGCT-P | KRAS-(WT)-FW/ KRAS-(M)-FW | Fig. S4-S5 |
| KRAS-RV-10C | P-TCCTACGCCCCCAGCT-P | KRAS-(WT)-FW/KRAS-(M)-FW | Fig. S4-S5 |
| KRAS-RV-11A | P-TCTACGCCACAAGCTC-P | KRAS-(WT)-FW/KRAS-(M)-FW | Fig. 2-5/Fig. S5, S14, S19, S20 |
| KRAS-RV-11T | P-TCTACGCCACTAGCTC-P | KRAS-(WT)-FW/KRAS-(M)-FW | Fig. S5 |
| KRAS-RV-11G | P-TCTACGCCACGAGCTC-P | KRAS-(WT)-FW/KRAS-(M)-FW | Fig. S5 |
| KRAS-FW-10A | P-TTTGGAGCTAGTGGCG-P | KRAS-(WT)-RV/KRAS-(M)-RV | Fig. 2-5/Fig. S5, S14, S19, S20 |
| KRAS-FW-10T | P-TTTGGAGCTTGTGGCG-P | KRAS-(WT)-RV/KRAS-(M)-RV | Fig. S5 |
| KRAS-FW-10C | P-TTTGGAGCTCGTGGCG-P | KRAS-(WT)-RV/KRAS-(M)-RV | Fig. S5 |
| KRAS-FW-11A | P-TTGGAGCTGGAGGCGT-P | KRAS-(WT)-RV/KRAS-(M)-RV | Fig. S5 |
| KRAS-FW-11G | P-TTGGAGCTGGGGGCGT-P | KRAS-(WT)-RV/KRAS-(M)-RV | Fig. S5 |
| KRAS-FW-11C | P-TTGGAGCTGGCGGCGT-P | KRAS-(WT)-RV/KRAS-(M)-RV | Fig. S5 |
| PIK3CA-RV-10A | P-TTCTCCTGCACAGTGA-P | PIK3CA-(WT)-FW/PIK3CA-(M)-FW | Fig. S6 |
| PIK3CA-RV-10G | P-TTCTCCTGCGCAGTGA-P | PIK3CA-(WT)-FW/PIK3CA-(M)-FW | Fig. 5/Fig. S6, S13, S16, S19, S20 |
| PIK3CA-RV-10C | P-TTCTCCTGCCCAGTGA-P | PIK3CA-(WT)-FW/PIK3CA-(M)-FW | Fig. S6 |
| PIK3CA-RV-11T | P-TCTCCTGCTCTGTGAT-P | PIK3CA-(WT)-FW/PIK3CA-(M)-FW | Fig. S6 |
| PIK3CA-RV-11G | P-TCTCCTGCTCGGTGAT-P | PIK3CA-(WT)-FW/PIK3CA-(M)-FW | Fig. S6 |
| PIK3CA-RV-11C | P-TCTCCTGCTCCGTGAT-P | PIK3CA-(WT)-FW/PIK3CA-(M)-FW | Fig. S6 |
| PIK3CA-FW-10A | P-TGAAATCACAGAGCAG-P | PIK3CA-(WT)-RV/PIK3CA-(M)-RV | Fig. S6 |
| PIK3CA-FW-10G | P-TGAAATCACGGAGCAG-P | PIK3CA-(WT)-RV/PIK3CA-(M)-RV | Fig. S6 |
| PIK3CA-FW-10C | P-TGAAATCACCGAGCAG-P | PIK3CA-(WT)-RV/PIK3CA-(M)-RV | Fig. 5/Fig. S6, S13, S16, S19, S20 |
| PIK3CA-FW-11T | P-TAAATCACTGTGCAGG-P | PIK3CA-(WT)-RV/PIK3CA-(M)-RV | Fig. S6 |
| PIK3CA-FW-11G | P-TAAATCACTGGGCAGG-P | PIK3CA-(WT)-RV/PIK3CA-(M)-RV | Fig. S6 |
| PIK3CA-FW-11C | P-TAAATCACTGCGCAGG-P | PIK3CA-(WT)-RV/PIK3CA-(M)-RV | Fig. S6 |
| EGFR-RV (Del-di-P) | P-TTTGCTTCTCTTAATT-P | EGFR-(WT)-FW-Del/EGFR-(M)-FW-Del | Fig. 5/Fig. S7, S13, S16, S19, S20 |
| EGFR-FW (Del-di-P) | P-TAAGGAATTAAGAGAA-P | EGFR-(WT)-RV-Del/ EGFR-(M)-RV-Del | Fig. 5/Fig. S7, S13, S16, S19, S20 |

**Table S2. ssDNA targets used in this study.**

| Name | Sequence | Presence |
|---|---|---|
| KRAS-(WT)- RV | TAGCTGTATCGTCAAGGCACTCTTGCCTAC | Fig. 2/Fig. S1, S3-S5 |
| | GCCACCAGCTCCAACTACCACAAGTTTATA | |
| KRAS-(WT)- FW | TATAAACTTGTGGTAGTTGGAGCTGGTGGC | Fig. 2/Fig. S5 |
| | GTAGGCAAGAGTGCCTTGACGATACAGCTA | |
| KRAS-(M)- FW | TATAAACTTGTGGTAGTTGGAGCTGATGGC | Fig. 2/Fig. S5 |
| | GTAGGCAAGAGTGCCTTGACGATACAGCTA | |
| KRAS-(M)- RV | TAGCTGTATCGTCAAGGCACTCTTGCCTAC | Fig. 2/Fig. S3-S5 |
| | GCCATCAGCTCCAACTACCACAAGTTTATA | |
| PIK3CA-(WT)-FW | AATTTCTACACGAGATCCTCTCTCTGAAATC | Fig. S6 |
| | ACTGAGCAGGAGAAAGATTTTCTATGGAG | |
| PIK3CA-(WT)-RV | CTCCATAGAAAATCTTTCTCCTGCTCAGTGA | Fig. S6 |
| | TTTCAGAGAGAGGATCTCGTGTAGAAATT | |
| PIK3CA-(M)-FW | AATTTCTACACGAGATCCTCTCTCTGAAATC | Fig. S6 |
| | ACTAAGCAGGAGAAAGATTTTCTATGGAG | |
| PIK3CA-(M)-RV | CTCCATAGAAAATCTTTCTCCTGCTTAGTGA | Fig. S6 |
| | TTTCAGAGAGAGGATCTCGTGTAGAAATT | |
| EGFR-(WT)-FW-Del | AAAATTCCCGTCGCTATCAAGGAATTAAGA | Fig. S7 |
| | GAAGCAACATCTCCGAAAGCCAACAAGGAA | |
| EGFR-(WT)-RV-Del | TTCCTTGTTGGCTTTCGGAGATGTTGCTTC | Fig. S7 |
| | TCTTAATTCCTTGATAGCGACGGGAATTTT | |
| EGFR-(M)-FW-Del | AAAGTTAAAATTCCCGTCGCTATCAAAACA | Fig. S7 |
| | TCTCCGAAAGCCAACAAGGAAATCCTCGAT | |
| EGFR-(M)-RV-Del | ATCGAGGATTTCCTTGTTGGCTTTCGGAGA | Fig. S7 |
| | TGTTTTGATAGCGACGGGAATTTTAACTTT | |

**Table S3. Primers used in this study.**

| Name | Target | Primer sequence (5'→3') | Presence |
|---|---|---|---|
| KRAS-158FW | KRAS (G12D) | GTGACATGTTCTAATATAGTC | Fig. 3-5/Fig. S8-S11, S14, S19, S20 |
| KRAS-158RV | KRAS (G12D) | GGATCATATTCGTCCACAAA | Fig. 3-5/Fig. S8-S11, S14, S19, S20 |
| PIK3CA-139FW | PIK3CA (E545K) | GAGACAATGAATTAAGGGAA | Fig. 5/Fig. S13, S16, S19, S20 |
| PIK3CA-139RV | PIK3CA (E545K) | GAAACAGAGAATCTCCATT | Fig. 5/Fig. S13, S16, S19, S20 |
| EGFR-157FW | EGFR (delE746-A750) | CTGTCATAGGGACTCTGGAT | Fig. 5/Fig. S13, S16, S19, S20 |
| EGFR-157RV | EGFR (delE746-A750) | GCCTGAGGTTCAGAGCCAT | Fig. 5/Fig. S13, S16, S19, S20 |
| M13F (-47) | 626 bp dsDNA target | CGCCAGGGTTTTCCCAGTCACGAC | Fig. 2b/Fig. S2, S5e, S6e, S7b |
| M13R (-48) | 626 bp dsDNA target | AGCGGATAACAATTTCACACAGGA | Fig. 2b/Fig. S2, S5e, S6e, S7b |

### Table S4. TaqMan probes and primers used in this study.

| Name | Sequence (5'-3') |
|---|---|
| PIK3CA-Forward Primer | 5’-GAACAGCTCAAAGCAATTTCTACAC-3’ |
| PIK3CA-Reverse Primer | 5’-AGCACTTACCTGTGACTCCATAG-3’ |
| PIK3CA (wt)-TaqMan Probe | 5’- CTGAAATCACTGAGCAGGA-3’ (FAM-BHQ1) |
| PIK3CA E545K-TaqMan Probe | 5’- TCTGAAATCACTAAGCAGGA-3’(VIC-BHQ1) |
| KRAS-Forward Primer | 5’-AGGCCTGCTGAAAATGACTG-3’ |
| KRAS-Reverse Primer | 5’-GCTGTATCGTCAAGGCACTCT-3’ |
| KRAS (wt)-TaqMan Probe | 5’-TTGGAGCTGGTGGCGTA-3’(VIC-BHQ1) |
| KRAS G12D-TaqMan Probe | 5’-TTGGAGCTGATGGCGTA-3’(FAM-BHQ1) |
| EGFR-Forward Primer | 5’- CCAGAAGGTGAGAAAGTTA-3’ |
| EGFR-Reverse Primer | 5’-TCGAGGATTTCCTTGTTG-3’ |
| EGFR (wt)-TaqMan Probe | 5’-CTTCTCTTAATTCCTTGATAGCGACGG-3’(FAM-BHQ1) |
| EGFR del E746-A750-TaqMan Probe | 5’-CGCTATCAAAACATCTCCGAAAGCC-3’(VIC-BHQ1) |

### Table S5. Clinical samples used in this study.

| Patient number | Cancer types | Genotype and mutation fraction* | Presence |
|---|---|---|---|
| Tissue sample 1 | Lung adenocarcinoma | *KRAS* G12D | Fig. 4b/Fig. S17 |
| Tissue sample 2 | Colorectal cancer | *KRAS* G12D | Fig. 4b/Fig. S17 |
| Tissue sample 3 | Endometrial cancer | *KRAS* G12D | Fig. 4b/Fig. S17 |
| Blood sample 1 | Colorectal cancer | *KRAS* G12D | Fig. 4b/Fig. S17 |
| Blood sample 2 | Colorectal cancer | *KRAS* G12D | Fig. 4b/Fig. S17 |
| Blood sample 3 | Colorectal cancer | *KRAS* G12D | Fig. 4b/Fig. S17 |
| Healthy donor 1 | ND** | — | Fig. 4b/Fig. S17 |
| Healthy donor 1 | ND** | — | Fig. 4b/Fig. S17 |

*Samples were analyzed with standard NGS protocol; **Not detected, possibly due to mutant allele not present.
